# Reliable sensory processing in mouse visual cortex through inhibitory interactions between Somatostatin and Parvalbumin interneurons

**DOI:** 10.1101/187062

**Authors:** Rajeev V. Rikhye, Ming Hu, Murat Yildirim, Mriganka Sur

## Abstract

Cortical neurons often respond to identical sensory stimuli with large variability. However, under certain conditions, the same neurons can also respond highly reliably. The circuit mechanisms that contribute to this modulation, and their influence on behavior remains unknown. Here we used novel double transgenic mice, dual-wavelength calcium imaging and temporally selective optical perturbation to identify an inhibitory neural circuit in visual cortex that can modulate the reliability of pyramidal neurons to naturalistic visual stimuli. Our results, supported by computational models, suggest that somatostatin interneurons (SST-INs) increase pyramidal neuron reliability by suppressing parvalbumin interneurons (PV-INs) via the inhibitory SST→PV circuit. Using a novel movie classification task, we further show that, by reducing variability, activating SST-INs can improve the ability of mice to discriminate between ambiguous stimuli. Together, these findings reveal a novel role of the SST→PV circuit in modulating the fidelity of neural coding critical for visual perception.

## INTRODUCTION

A longstanding aim of systems neuroscience is to relate neural activity to perception. In visual perception, it has been established that a key role of the primary visual cortex (V1) is to transform raw sensory information from the environment into a low-level percept^1^. Surprisingly, under laboratory conditions, V1 pyramidal neurons respond to repetitions of identical sensory stimuli with spike trains that vary greatly in both the number and the timing of spikes^2,3^. Although it is known that this unreliability limits stimulus selectivity^4^, its impact on perception, and more specifically, visually-guided behavior, remains unknown. Notably, a large part of this variability is generated internally within the cortex, as the same neuron can respond either reliably or unreliably under different conditions^5,6^. For example, increasing the size of the stimulus within the receptive field or its statistics from simple (gratings) to complex (natural scenes) increases reliabilty^7^. Additionally, arousal and attention decreases variability and increases task performance^8–11^. Collectively, these findings suggest that mechanisms might exist within the cortex to modulate response reliability depending on processing demands^12^. The goal of this study is to elucidate these mechanisms and study their influence on visual behavior in mice.

Inhibitory neurons (INs) play an important role in controlling cortical activity at various temporal and spatial scales^13^. Hence, changes in cortical inhibition might be a potential mechanism responsible for modulating response reliability. It has been noted that inhibitory post synaptic potentials measured in cortical pyramidal neurons are precisely delayed relative to excitatory potentials during epochs of reliable firing^14^. This delayed inhibitory input is believed to quench stochastic excitatory inputs by limiting integration to a small window during which reliable spiking can occur^15^. Moreover, chronically blocking inhibition sharply decreases response reliability^16,17^. However, given their computationally diversity^18^, the specific role of different IN subtypes in modulating response reliability remains poorly understood. This study focuses on the two major IN classes – parvalbumin-(PV) and somatostatin-expressing (SST) INs – which provide distinct inhibitory control over pyramidal (EXC) neurons in layer 2/3 of mouse V1. PV-INs provide rapid, shunting inhibition onto the somatic compartment of EXC neurons^19^, and as a consequence, are able to powerfully control the response gain^20,21^ and spike timing^22^ of their targets. SST-INs, on the other hand, inhibit the distal dendrites of pyramidal neurons, where they can control synaptic integration^23–25^. SST-INs also receive strong recurrent excitation and have been found to influence network integration^26,27^. Importantly, these INs do not act independently as SST-INs also inhibit PV-INs^28,29^. Through this inhibitory SST→PV circuit, SST-INs have the ability to control the inhibitory tone of both the dendritic and the somatic compartments, making them ideal candidates to modulate variability both at the level of synaptic input and spiking output. However, little evidence exists to support this hypothesis.

Here we developed a new line of double transgenic mice, and used a multifaceted approach to study how interactions between PV and SST-INs contribute to reliable sensory processing in V1. Using dual-wavelength calcium imaging, we found that SST-INs were more active during epochs of reliable pyramidal cell firing whereas PV-INs were more active during epochs of unreliable firing. This complementary activity was due to the inhibitory SST→PV circuit. Using temporally-limited optical perturbations and computational models, we found that SST-INs improve reliability by suppressing PV-INs. Activating SST-INs as mice performed a natural movie classification task improved discrimination performance, whereas activating PV-INs had the opposite effect, demonstrating that variability in V1 is detrimental to visual perception. Thus, our work identifies a novel mechanism in which PV and SST-INs work cooperatively, via the SST→PV circuit, to modulate the fidelity of sensory processing.

## RESULTS

### SST and PV-INs have mutually exclusive dynamics during epochs of reliable firing

We measured the reliability of V1 excitatory pyramidal (EXC) neurons in awake, passive mice to repeated presentations of naturalistic movies (five different movies) using two-photon calcium imaging. We used natural movies because they are known to drive sparse and reliable responses from EXC neurons across different species^30–33^. To target EXC neurons in layer 2/3 of V1, we expressed the genetically encoded calcium indicator (GECI) GCaMP6f in PV-tdTomato and SST-tdTomato mice (PV-Cre and SST-Cre x Ai14) via stereotactic injections of an adeno-associated virus (Fig. 1a, also see Methods). Since PV and SST-INs in these mice express the red fluorescent protein, tdTomato, we reasoned that the majority of tdTomato-negative neurons would be EXC neurons. We quantified the trial-to-trial response reliability of these neurons for each movie by computing the average of all pair-wise correlations (corrected for differences in the mean firing rate) between single trial responses (see Methods). By this definition, reliability measures the degree of trial-to-trial similarity in evoked responses to a given movie. Expectedly, response reliability was strongly negatively correlated with between-trial variability (Fig. 1b, c), as the least variable neurons also had responses that were highly similar across trials.

**Figure 1.**
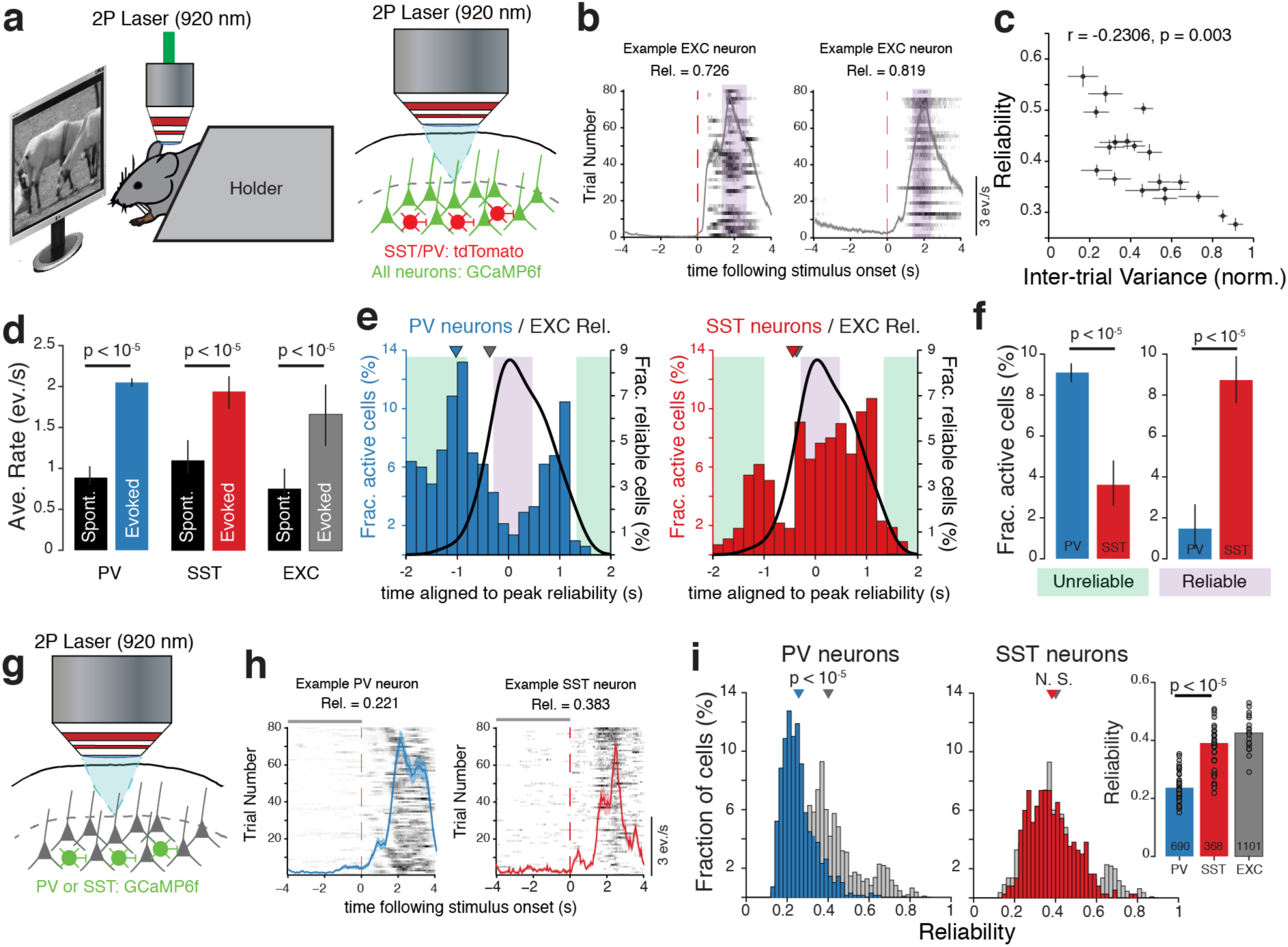
SST and PV-INs respond during distinct epochs of EXC neuron activity. **(a)** Schematic showing experimental setup and method to record from EXC neurons. **(b)** Raster plots (trials vs. time) of two simultaneously recorded EXC neurons showing reliable and sparse responses to the same movie. Gray lines show trial-averaged responses and shaded areas denote SEM over trials. Shaded purple bar shows time period (epoch) when these EXC neurons are reliably activated. **(c)** Scatter plot showing strong negative correlation between inter-trial variance and response reliability. Each data point is the mean response reliability and the across-trial variance of each imaged population, error bars are SEM (19, each with 22-87 neurons, 10 mice). **(d)** Comparison between evoked (averaged from 5 different movies) and spontaneous (spont., gray screen) activity for all the three cell types, expressed in number of inferred events per second. All cell types showed a significant increase in evoked response rate compared to spontaneous activity. Errorbars, SEM. **(e)** Histogram showing the fraction of active PV (left) and SST-INs (right) in 200 ms time bins aligned to peak EXC population reliability. Triangles above the histograms indicate mean time to peak activity. Significant difference between PV and EXC neuron activation times (p < 10^−6^) but no significant difference between activation times for SST and EXC neurons (p = 0.129). **(f)** Bar plots comparing the median fraction of active PV and SST-INs during epoch of unreliable and reliable EXC neuron firing respectively. Errorbars, 95% CI. Data in H and I are from 10 mice (1101 EXC neurons, 120 SST-INs, 186 PV-INs). **(g)** Method to image INs. **(h)** Example raster plot of a PV and a SST-IN to the same movie. Format same as (b). **(i)** Histogram of PV and SST-IN reliability in relation to EXC neuron reliability (gray). Triangles above the histograms indicate mean reliability pooled over all neurons. Inset compares median reliability for all cell types. Each data point is the median reliability of each imaged population. Data from: PV = 8 mice (690 neurons); SST = 8 mice (368 neurons); EXC = 10 mice (1101 neurons). All p-values computed using grouped, Bonferroni-corrected rank-sum test.

In all imaged populations (10 mice, 1101 neurons), EXC activity patterns spanned a range from highly stereotyped and reliable responses to weak and variable activity. On average, however, most neurons (37.7±15.4%) in each population responded reliably (>0.4, i.e. similar responses on at least 40% of the trials) to at least one movie. EXC neuron activity was typically punctuated by brief epochs of highly reliable responses (see examples in Fig. 1b). The duration and magnitude these epochs varied from movie-to-movie, and were due to differences in the spatiotemporal statistics of these movies (Supplementary Fig. 1a-d). These observations agree with previous studies^32,33^, and establish that naturalistic movies can drive EXC neurons in mouse V1 to respond reliably and with low variability between trials.

With the aim of elucidating the inhibitory mechanisms responsible for this reliable coding of naturalistic scenes, we first quantified the response properties of different IN subtypes to the same movies. Similar to EXC neurons, both PV and SST-INs responded to these movies with an approximate two-fold increase in response rate over spontaneous activity (Fig. 1d). Notably, there was no significant difference in evoked response rates between these INs (p = 0.131, rank-sum test). Additionally, movies, which activated a greater number of reliably responding EXC neurons, also recruited a comparable fraction of INs (Supplementary Fig. 1a). These results suggest that the same feed-forward factors, such as stimulus properties, that drive EXC neurons to fire reliably are also effective at recruiting both PV and SST-INs (Supplementary Fig. 1f, g).

To further examine the relationship between EXC reliability and IN activity, we characterized PV and SST-IN activity around epochs of reliable/unreliable EXC neuron firing. In each simultaneously recorded neural population, we computed the reliability of EXC neurons and the mean rate of PV/SST-INs in 200 ms time bins following stimulus onset. In each time bin, we then computed the fraction of fraction of reliably responding EXC neurons (i.e. neurons with reliability >0.4), which gave us a measure of how consistently the population responded to each movie repetition, and the fraction of active PV/SST-INs. Aligning the fraction of active INs to the epoch of maximum reliability (which facilitated comparisons between different movies and populations, that each had different response dynamics), revealed that the majority of PV-INs were active during epochs of unreliable EXC neuron firing (Fig. 1e, f). In contrast, SST-INs were most active during epochs when EXC neurons were most reliable. Therefore, although similar in response magnitude, PV and SST-INs are active during distinct epochs of EXC neuron activity.

Do INs also respond reliably to these movies? To better quantify the reliability of the different IN subtypes, we restricted GECI expression to INs by injecting an adeno-associated virus encoding a Cre-dependent variant of GCaMP6f in either PV-Cre or SST-Cre mice (Fig. 1g). This method allowed us to avoid neuropil contamination from neighboring EXC neurons. Interestingly, although PV-INs responded strongly to most movies, their responses were much more variable between trials (Fig. 1h, i). As a consequence, PV-INs were less reliable than EXC neurons. In contrast, SST and EXC neurons had similar reliability values (Fig. 1h, i).

Taken together, these results suggest two complementary modes of inhibition with SST-INs providing reliable inhibition during epochs of reliable EXC firing and PV-INs providing unreliable inhibition during unreliable epochs.

### The joint dynamics of SST and PV-INs scale with EXC reliability

Given this complementary relationship between PV and SST-INs, we next sought to characterize the trial-by-trial interactions between PV and SST-INs within the same neuronal population. To gain independent genetic access to both cell types^34^, we crossed SST-Cre mice with PV-FlpO mice to create a new strain of double transgenic mice (referred to as SXP mice), where SST-INs express Cre recombinase and PV-INs express flp recombinase. To monitor the joint activity of these IN subtypes *in vivo*, we concurrently expressed the red GECI (jRGECO1a) in SST-INs and the green GECI (GCaMP6f) in PV-INs in SXP mice and performed dual-wavelength calcium imaging (Fig. 2a). Using custom optics, we scanned the same field-of-view with two mutliplexed lasers – one tuned to 1020nm to excite jRGECO1a and the other tuned to 920nm to excite GCaMP6f (see Methods). These wavelengths optimally excite each GECI with very little spectral overlap^35^ (Supplementary Fig. 2a). Notably, we observed a negligible fraction of co-labeled cells (data not shown), confirming that the labeled PV and SST-INs were indeed non-overlapping cell types^18^.

**Figure 2.**
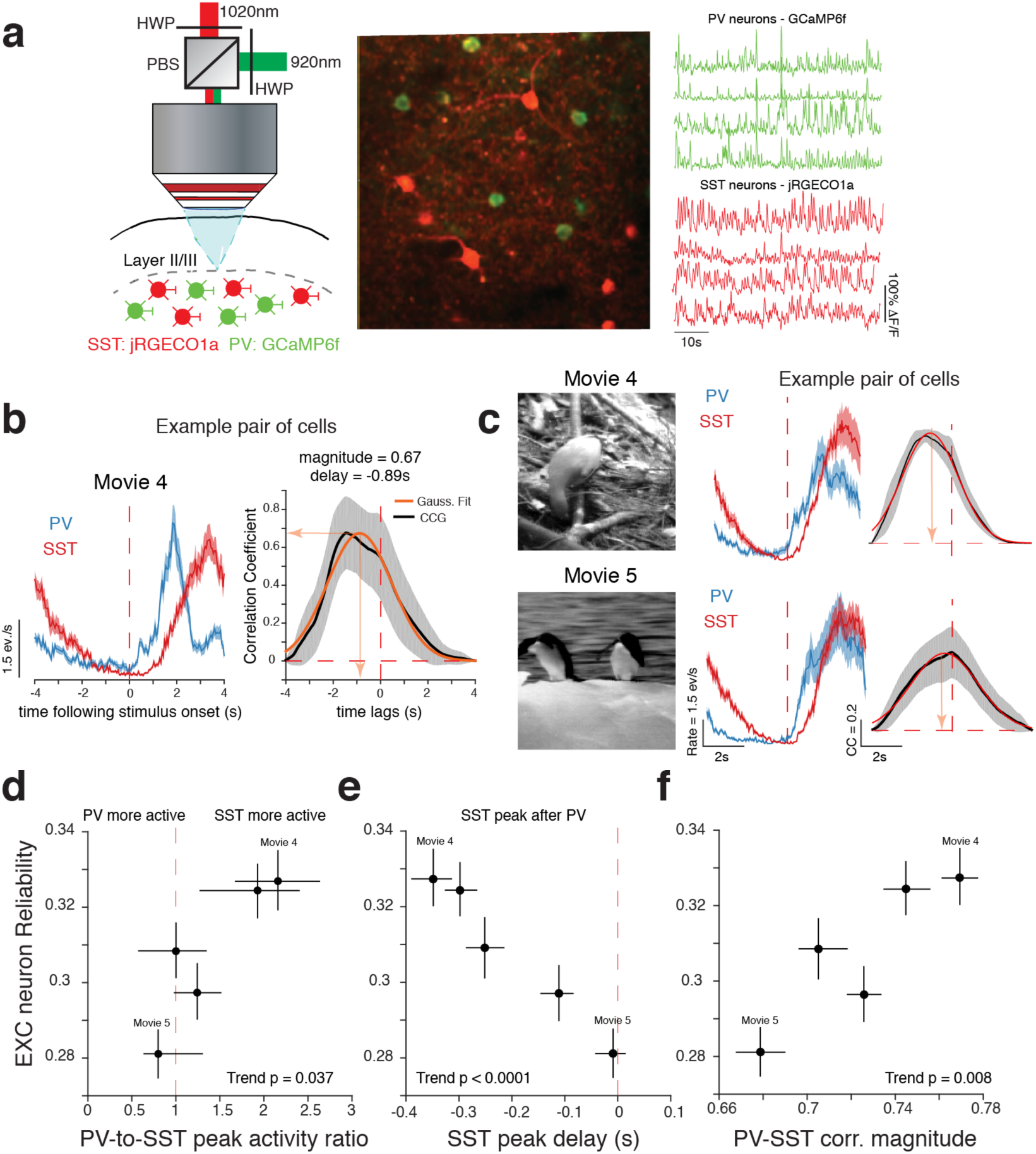
SST-INs are temporally delayed relative to PV-INs in reliably processed movies. **(a)** Left: Experimental setup. Briefly, a 1020nm laser and a 920nm laser were combined using a half wave plate (HWP) and a polarizing beam splitter (PBS) to optimally activate jRGECO1a and GCaMP6f in SST and PV-INs respectively (see Methods). Middle: Example field of view showing co-labeled PV and SST-INs. Image covers a cortical area of 150μm x150μm. Right: Example calcium transients from simultaneously recorded interneurons. **(b)** Left: Trial-averaged responses from a pair of simultaneously recorded PV and SST-INs. Right: Cross-correlogram (CCG) of this pair. Orange line shows Gaussian fit to trial averaged CCG. Shaded areas, SEM over trials. **(c)** Difference in correlation for two different movies for the same PV and SST-IN pair. **(d-f)** More reliable movies have a stronger SST peak activity compared to PV **(d)**, longer delays between SST and PV peak activity **(e)** and stronger PV-SST correlation at peak delay **(f)**. Data are from 2292 pairs, 5 mice. Data points denote median ± 95% CI for each movie. P-values computed using F-test to measure significance of the trend relative to a constant model.

To quantify the temporal relationship between these INs we first computed cross-correlograms (CCG) single trial responses of PV and SST-INs. We then estimated the activation delay and the correlation strength between each pair from the Gaussian function that best fit the CCG (Ave. fit R^2^ = 78.9±4.6%, Fig. 2b). Across all recorded pairs, we measured an average time lag of −321 ms (CI: −360 to −283 ms), indicating that most SST-INs respond after PV-INs (75.44% of pairs, 5 mice, Fig. 2b, see also Supplementary Fig. 2b). Note that this delay measures the time difference between the peak calcium activity of SST and PV-INs, and not the spiking onset latency, which is significantly shorter^36^.

Interestingly, the strength and timing of these interactions also differed between movies; with some movies evoking more temporally correlated and delayed activity than others (Fig. 2c). Given that EXC neuron reliability also varied between movies, we next sought relate joint PV-SST activity with EXC reliability. However, due to technical limitations, we were unable to record from all three cell types simultaneously. Thus, we compared simultaneously recorded joint PV-SST activity to EXC neuron reliability obtained from separate mice but using the same movies, and at similar cortical locations (data from Fig. 1). Movies that were more reliably processed (i.e. those with higher median EXC neuron reliability) evoked stronger activity from SST than PV-INs (Fig. 2d), which was consistent with our data in Fig. 1h. Furthermore, PV-SST pairs were more strongly correlated and had longer delays in movies that were more reliably processed, than movies that were less reliably processed (Fig. 2e, f). Multivariate linear regression analysis confirmed that the ratio of PV-to-SST activity, lag duration and the correlation strength between PV and SST-INs were all significant predictors of EXC neuron reliability (p = 0.0038, F-test relative to constant model).

Our observation that peak SST activity is delayed relative to PV agrees with recent calcium imaging results showing that, across many different cortical areas, SST-INs respond after PV-INs^37^. Furthermore, in V1, sinusoidal gratings also elicit delayed responses^36^, albeit at a much shorter time scale. Possible mechanisms that could account for this delay include: (1) pooling of inputs from EXC neurons^38^, or (2) the inhibitory connection between SST and PV-INs^28^. We took advantage of SXP mice to provide two additional pieces of evidence to support the latter claim. First, silencing SST-INs optically with ArchT strongly increased PV-IN firing rate (Supplementary Fig. 3a), confirming a strong inhibitory connection between SST and PV-INs. Second, optically activating SST-INs with channelrhodopsin-2 (ChR2), suppressed PV-INs and increased the duration off the temporal delay between PV and SST-INs (Supplementary Fig. 3b). Therefore, these results suggest that this temporal delay is the result of the SST→PV circuit, which suppress PV-INs as SST activity ramps up.

Our experiments in SXP mice thus revealed the surprising result that the strength of temporal interactions between PV and SST-INs, which is coordinated by the SST→PV circuit, vary with EXC neuron reliability. Therefore, it is possible that both SST-INs and PV-INs might work together to reduce variability.

### Intact inhibition from both IN subtypes is necessary for reliable coding

If reliable coding requires intact cortical inhibition, then perturbing the activity of INs should reduce reliability. To verify this claim, we used designer receptors activated by designer drugs (DREADDs) to chronically, and non-specifically perturb PV and SST-INs, while monitoring EXC reliability using calcium imaging. In separate PV-and SST-Cre mice we expressed either the inhibitory (hM4D(Gi)) or the excitatory (hM3D(Gq)) DREADD variant (Supplementary Fig. 4a; see Methods). Intraperitoneal injection of clozapine-N-oxide (CNO) either increased or decreased IN activity in a chronic, non-specific manner (Supplementary Fig. 4b, f). Surprisingly, we found that either increasing or decreasing the activity of both PV and SST-INs strongly reduced EXC neuron reliability compared to saline controls (Supplementary Fig. 4c, g). This reduction in reliability was independent of the change in firing rate, as both suppressed and activated EXC neurons showed a reduction in reliability (Supplementary Fig. 4d, h). Notably, these perturbations disrupted the complementary relationship that these INs had with EXC reliability. Therefore, these chemogenetic manipulations not only demonstrate that intact inhibition from both IN subtypes is necessary for reliable coding, but also show that disrupting the precise relationship between PV and SST firing and EXC reliability can reduce reliability.

### SST and PV-INs bi-directionally modulate EXC neuron reliability

Using SST-Cre and PV-Cre mice crossed with the Cre-dependent ChR2 mice (Ai32), we next asked how photo-activating each IN subtype specifically during epochs of either reliable or unreliable firing affected reliability. Since the timing and duration of these epochs are heterogeneous within any given population, we had no way of estimating *a priori* when EXC neurons would respond reliably during a particular movie. To circumvent this issue, we developed an optical stimulation strategy to activate ChR2-expressing PV and SST-INs (PV-Cre and SST-Cre x Ai32) at 22 different time points that spanned the entire duration of a movie (Fig. 3a, see Methods). Specifically, we first created a distribution of stimulation events that spanned the duration of a movie, with the first and last events coincident with the onset and offset of the stimulus respectively, and the remaining events occurring at fixed interval of 200 ms. Then, on every movie repetition (trial), we picked a stimulation event from this distribution in a pseudo-randomized manner (as illustrated in Fig. 3a). These brief blue (473 nm) laser pulses were sufficient to reliably excite both PV and SST-INs at all time points during a movie (Supplementary Fig. 5). Thereafter, with *post hoc* analysis, we focused primarily on stimulation events that coincided with epochs of either reliable or unreliable firing.

**Figure 3.**
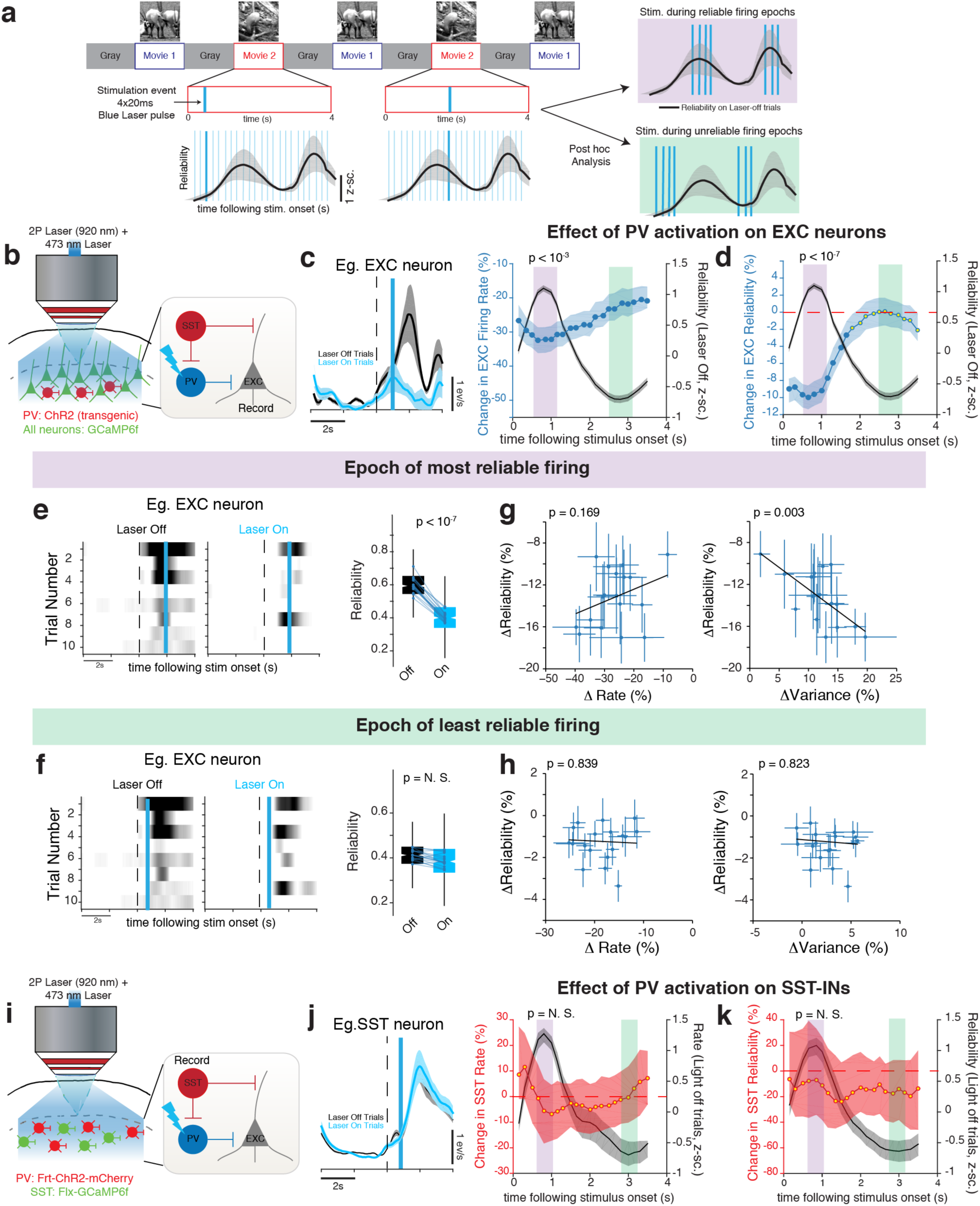
Increasing PV-IN activity reduces EXC neuron reliability. **(a)** Cartoon describing random stimulation strategy. A brief laser pulse (stimulation event) was applied at 22 equally spaced time points during a 4s movie (illustrated by the light blue lines). At each movie repetition, stimulation event time is drawn from this distribution at random (indicated by dark blue line). The bottom plots show the timing of each stimulation event in relation to the reliability of an example EXC neuron (black line). Following this, post hoc analysis was used to identify stimulation events that occurred within periods of reliable firing and unreliable firing (shaded purple and green respectively). **(b)** Cartoon of experimental setup. **(c)** Left: Representative example of an EXC neuron that is suppressed following PV activation. Blue line indicates the time of the stimulation event. Right: Change in firing rate for each PV stimulation event. To facilitate comparisons between movies and mice, all neurons were aligned to have a maximum reliability at 1s. All shaded areas are the 95% CI of the median. Yellow circles represent non-significant change (relative to 0) and were computed using a permutation test. P-values (Bonferroni-corrected rank-sum test) compare changes in firing rate between epochs of reliable vs. unreliable responses (shaded bars). **(d)** Change in EXC reliability for each stimulation event. **(e)** Left: Representative raster plot of an EXC neuron showing a reduction in reliability following PV activation during the reliable firing epoch. Right: Box-whisker plots summarizing the effect of PV activation on EXC neuron reliability. Each dot is pooled data from one population. P-value computed using Bonferroni-corrected Wilcoxon rank-sum test. **(f)** Same as **(e)**, but shows no change in reliability when PV-INs are activated during epochs of unreliable firing. **(g-h)** Scatter plots quantifying the relationship between ΔReliability and a change in rate (ΔRate, left) or a change in between-trial variability (ΔVariance, right). Error-bars are the 95% CI of the median. P-values computed from multivariate linear regression analysis. **(i)** Cartoon describing method to study the effect of PV activation on SST-INs. **(j)** Left: Representative SST-IN that shows no change following PV activation. Right: No change in SST rate for all PV activation events. **(k)** No change in SST reliability for all PV activation events.

We first investigated the effect of perturbing PV-IN activity (Fig. 3b). As expected, activating PV-INs suppressed EXC neurons shortly after laser onset. The magnitude of suppression was strongest when EXC neurons were most reliable and weakest when activity was unreliable (Fig. 3c). This suppression lasted for ~600 ms, reflecting polysynaptic effects caused by activating a large number of PV-INs, and we restricted our analysis of reliability during this time period of maximum suppression (Supplementary Fig. 7). Activating PV-INs significantly reduced the reliability of EXC neurons (Fig. 3d). In particular, increasing the strength of PV inhibition during epochs of reliable firing led to a ~20% reduction in EXC neuron reliability (Fig. 3e), whereas further increasing PV-IN activity during epochs of unreliable firing, when they are normally most active (Fig. 1h), did not change reliability (Fig. 3f).

These observed changes in reliability could be due to changes in either mean response rate (ΔRate) or between-trial variance (ΔVariance) or both. For example, a strong decrease in response rate, without a change in variability, could also reduce reliability. To quantify the effect these attributes had on ΔReliability, we used multivariate linear regression (model: ΔReliability ~ 1 + ΔRate + ΔVariance). Surprisingly, we did not observe a correlation between ΔRate and ΔReliability following PV-IN activation in either response epoch, as neurons that were suppressed more did not exhibit a larger decrease in reliability (Figs. 3g, h). Instead, the reduction in reliability following PV-IN activation during periods of reliable firing was strong correlated with an increase in firing rate variance between the trials (ΔVariance: p < 10^−3^ vs. ΔRate: p > 0.05, t-test). This implies that the change in reliability following PV-IN activation was due to an increase in variability rather than a change in rate.

Given the highly recurrent architecture of V1 layer 2/3, it is possible that, in addition to EXC neurons, perturbing PV-INs would also affect SST-INs. To answer this question, we conditionally expressed a FlpO-dependent ChR2 in PV-INs and a Cre-dependent GCaMP6f in SST-INs in SXP mice (Fig. 3i). Surprisingly, regardless of when in the movie we activated PV-INs, we did not observe a suppression of SST-INs (Fig. 3j) or a change of SST-IN reliability (Fig. 3k). This result implies that there is not an inhibitory PV→SST connection^28^, and the reduction of EXC neuron rate is not sufficient to alter SST-IN activity. Importantly, this result implies that the reduction in EXC neuron reliability following PV activation is not due to a change in SST→EXC inhibition, but rather due to direct PV→EXC inhibition.

In contrast to PV-INs, activating SST-INs had a much weaker suppressive effect on EXC neurons (Fig. 4a, b, p < 10^−4^, one-way Kruskal-Wallis ANOVA relative to PV-ChR2). This was in part due to a disinhibitory effect caused by a lifting of PV inhibition following SST activation as ~27% of the neurons (8/20 populations) showed an increased rate following SST activation. Also in contrast to PV activation, increasing the strength of SST inhibition increased the reliability of EXC neurons (Fig. 4c). This effect was most significant when SST-INs were activated during epochs of unreliable firing (Fig. 4e). This increase in reliability was correlated with a strong reduction in trial-to-trial variance and a modest increase in response rate (Fig. 4g). Multivariate linear regression confirmed that both variables had a significant effect on ΔReliability (ΔVariance: p < 10^−5^ and ΔRate: p < 10^−2^, t-test). Interestingly, further increasing the activity of SST-INs during periods of reliable firing did not alter EXC neuron reliability and only marginally reduced trial-to-trial variance (Figs. 4d, f).

**Figure 4.**
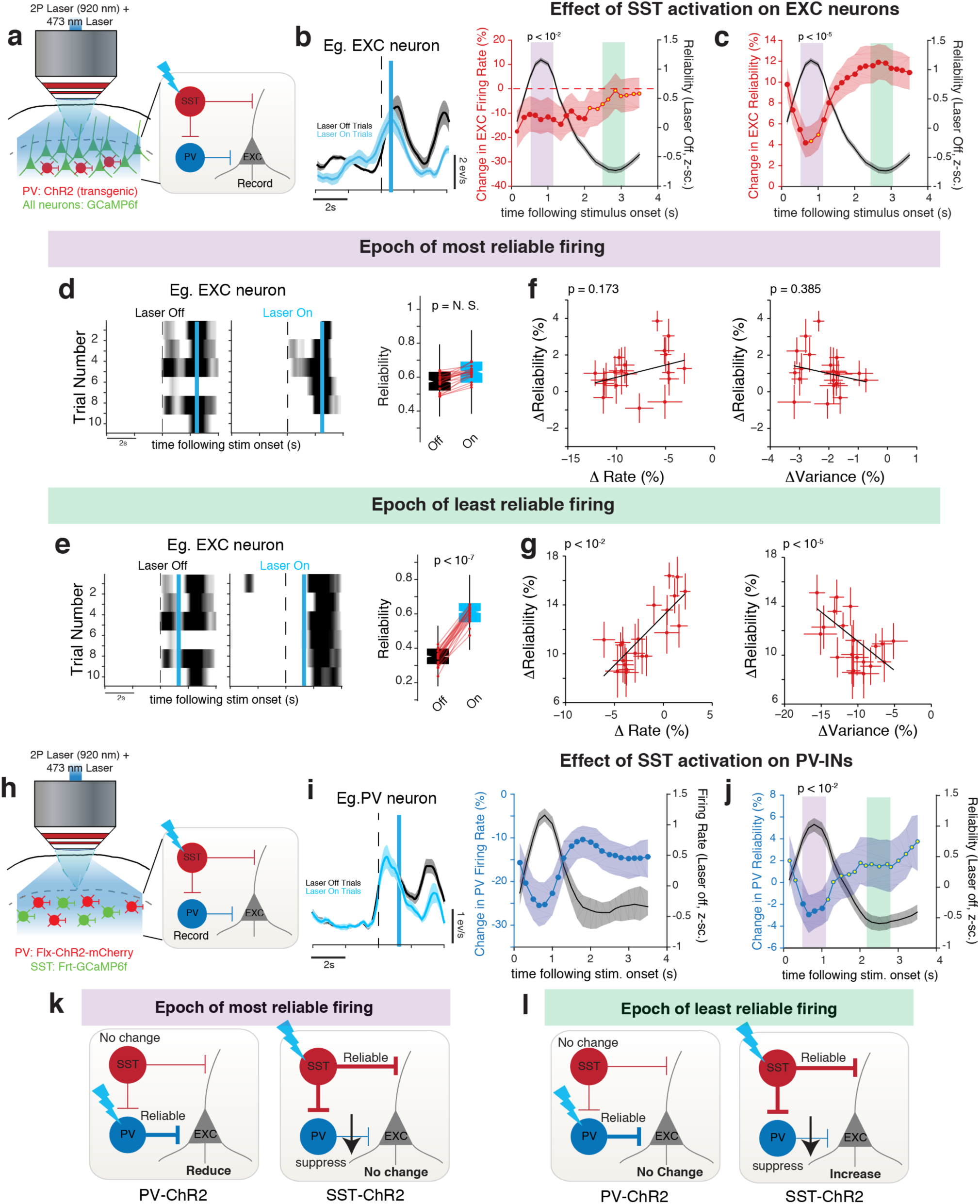
Increasing SST-IN activity increases EXC neuron reliability. **(a)** Cartoon of experimental setup. **(b)** Left: Representative EXC neuron that shows a modest decrease in firing rate following SST activation (denoted by blue bar). Right: Change in firing rate for each SST stimulation event shown in relation to EXC neuron reliability on light-off trials. **(c)** Change in EXC neuron reliability for each SST stimulation event. **(d)** Representative raster plot of an EXC neuron and Box-whisker plot showing no change in reliability following SST activation during epoch of most reliable firing. **(e)** Same as **(d)**, but showing that SST activation during epoch of least reliable firing can increase EXC neuron reliability. **(f-g)** Scatter plots quantifying the relationship between ΔReliability and a change in rate (ΔRate, left) or a change in between-trial variability (ΔVariance, right). Error-bars are the 95% CI of the median. P-values computed from multivariate linear regression analysis. Data in **(b-g)** are from 8 SST-ChR2 mice (622 neurons, 19 populations). **(h)** Cartoon describing method to study the SST activation on PV-INs. **(i)** Left: Representative PV-IN that is suppressed following SST activation. Right: PV-IN firing rate is significantly suppressed for all SST activation events. **(j)** SST activation reduces the reliability of PV-INs. Data in same format as Fig. 3 and are from 4 SXP mice (372 PV neurons) **(k-l)** Cartoon summarizing photo-activation results from Figs. 3 and 4.

Using SXP mice, we found that SST activation strongly suppressed PV-INs, regardless of when the stimulation occurred during the movie (Fig. 4h, i). Surprisingly, unlike EXC neurons, activating SST-INs when PV-INs were at their least reliable did not change their reliability (Fig. 4j). Instead, PV-INs became more unreliable. Therefore, increasing SST inhibition influences both EXC neuron reliability and the dynamics of PV-IN inhibition.

As a control, we repeated these activation experiments in mice, which expressed the red fluorescent protein tdTomato in either PV or SST-INs, instead of ChR2. In these mice, we found no significant change in either response rate or reliability of EXC neurons following laser stimulation (Supplementary Fig. 6). Additionally, we found that inferring firing rates via deconvolution did not influence our reliability calculation, as neurons that had reliable calcium transients also had reliable inferred rates (Supplementary Fig. 7a, b).

Altogether, these results demonstrate complementary roles of PV and SST-INs in modulating EXC neuron reliability (summarized in Fig. 4k, l). In particular, increasing PV inhibition when EXC neurons are reliable decreases their reliability by increasing variability between trials, but does not alter SST-IN dynamics. On the other hand, increasing SST inhibition when EXC neurons are unreliable increases their reliability by a combined effect of decreased variability, and to a lesser extent, a disinhibitory increase in response rate, caused by a suppression of PV-INs.

### Computational model predicts that SST-INs increase reliability by suppressing PV-INs

How do SST-INs increase EXC neuron reliability? Given the complementary relationship between PV and SST-INs, we hypothesized that the inhibitory SST→PV circuit might play a role in coordinating activity between these INs and consequently modulating EXC neuron reliability. To test this hypothesis, we developed a computational model of V1 microcircuit dynamics that simulated the mean firing rate of different neural subtypes (Fig. 5a, see Methods). Our model comprised four rate-based units (EXC, SST, PV, and VIP-INs) that were interconnected through connectivity parameters^39^. Although, we did not investigate VIP-INs experimentally, we included them in the model for completeness. EXC, SST and PV units in our model received “visual” input from a bank of linear-nonlinear-Poisson (LNP) units^40^, each with Gabor-like spatiotemporal receptive fields of different orientations and spatial frequencies (Supplementary Fig. 8, see Methods). Due to the stochastic nature of the Poisson process, each trial produced “visual” inputs, that differed in both the number and timing of spikes. As previously reported, this method allowed us to accurately capture both the temporal dynamics and reliability of the same movies that we used in our experiments^7^. Additionally, each unit in the model also received independent stochastic Poisson noise, which modeled background inputs. As a result, each unit, except for VIP, had an independent (uncorrelated) and a shared source of variability.

**Figure 5.**
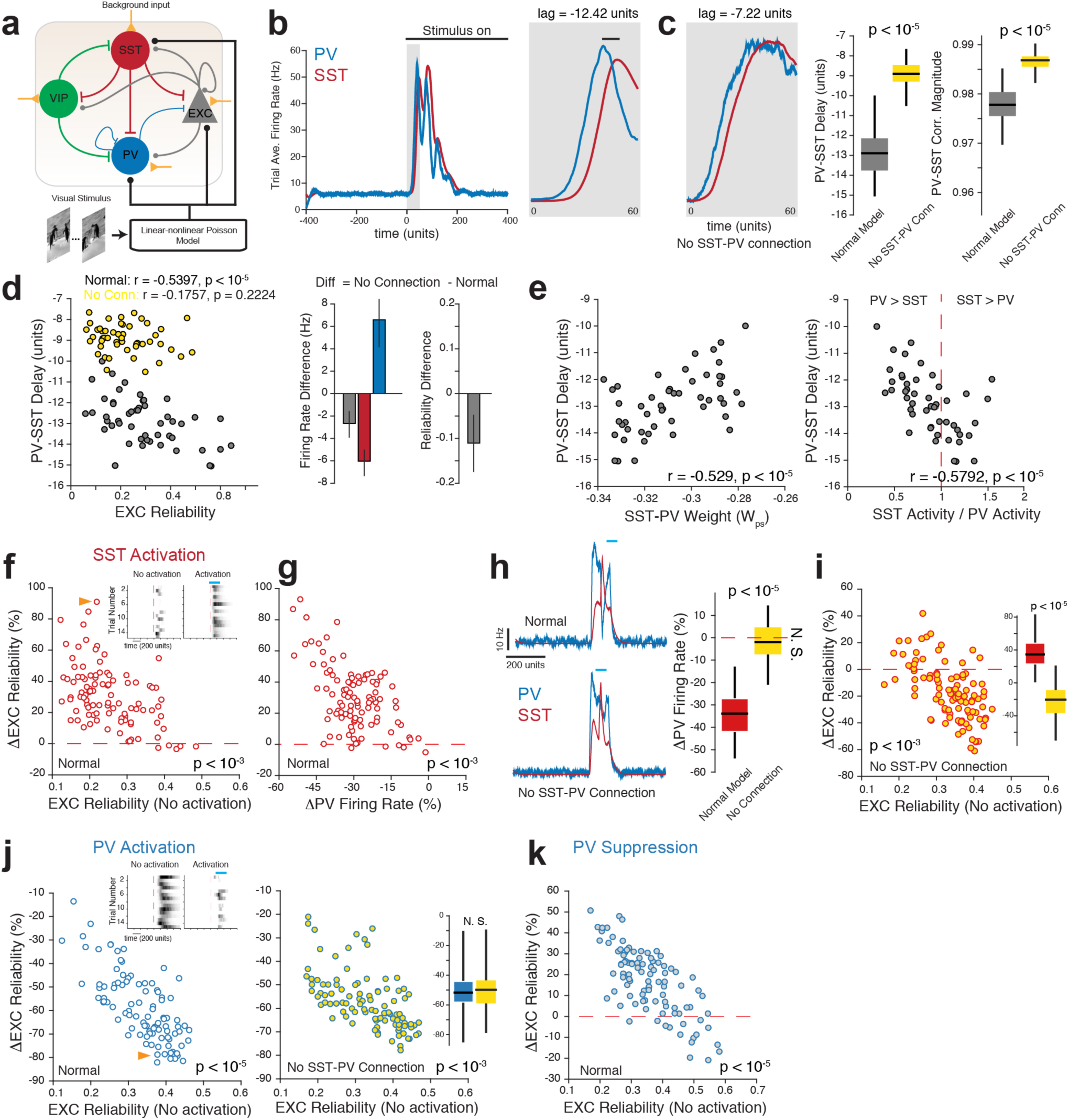
Computational model predicts that SST-INs increase reliability by suppressing PV-INs. **(a)** Cartoon illustrating connectivity between the four major units simulated in this model. Round connections indicate excitatory synapses, while blunt connections indicate inhibitory synapses. see Methods for details of the linear-nonlinear Poisson model. **(b)** Representative simulation showing the response of PV and SST units to a natural movie. Inset shows a zoomed view of the onset dynamics to highlight the temporal lag between PV and SST units. **(c)** The delay between PV and SST units is reduced when the SST→PV connection is removed. Box-whisker plots quantify the change in time lag and correlation strength between PV and SST units with and without the SST→PV connection. Data is pooled from 100 simulations each with randomly drawn connection weights. P-value computed using Kruskal-Wallis one-way ANOVA. **(d)** Left: Significant correlation between PV-SST delay duration and EXC unit reliability for the normal model, which is lost when SST→PV connection is cut. Middle and Right: Removing the SST→PV circuit increases PV unit firing rate while suppressing SST and EXC units. This perturbation also decreases reliability. Error bar, SEM over simulations. **(e)** Regression plots showing that the both the weight of the SST→PV connection and the SST-to-PV activity ratio is predictive of the delay duration. **(f)** Model predicts that activating SST activation during epoch of unreliable firing will increase the reliability of EXC units. Inset, shows representative raster plot of an EXC unit (indicated by orange arrow). **(g)** Large changes in EXC unit reliability are associated with a large decrease in PV unit firing rate. **(h)** SST-induced suppression is reduced when the SST→PV connection is cut. **(i)** Without the SST→PV circuit, our model predicts that SST activation will reduce the reliability of EXC neurons. Inset, Box-whisker plots comparing the change in reliability with and without an intact SST→PV connection. **(j)** Model predicts that PV activation will reduce reliability (left) and removing the SST→PV connection will not affect this change in variability. **(k)** Suppressing PV units will result in an increase in variability. All data points are an independent simulation in which a natural movie is repeated 30 times. To test robustness, we repeated each simulation 100 times, each with randomly drawn connection weights. P-values in the scatter plots are computed from linear regression (see Methods).

First, we asked whether this model could recapitulate and explain the relationship between PV-SST delay and EXC reliability (Fig. 2). As in our experimental data, SST units in our model also lagged behind PV units with a variable delay (Fig. 5b). Removing the SST→PV connection reduced the duration of the lag, and unexpectedly increased PV-SST correlation strength (Fig. 5c). This further confirms that the temporal relationship observed between PV-SST pairs *in vivo* is due to the inhibitory SST→PV circuit. Also similar to our experimental results, we observed a correlation between the duration of the PV-SST lag and EXC reliability, such that models with higher EXC reliability also had more delayed SST peak activity relative to PV peak activity (Fig. 5d). Removing the SST→PV connection abolished this relationship, increased PV unit activity and reduced EXC neuron reliability (Fig. 5d). We performed multivariate regression analysis to identify which variables contributed most to this temporal relationship. Interestingly, the strength of the SST→PV connection and the activity fraction of SST units to PV units were the biggest predictors of the lag duration (Fig. 5e). This implies that conditions, which strongly recruit SST-INs, such as reliable EXC neuron firing, will increase the dynamics of joint PV-SST activity. In further support of this mechanism, our model also accurately predicts that perturbing PV and SST activity for the entire stimulus duration will jointly reduce reliability (Supplementary Fig. 4e, i). Therefore, intact joint PV-SST dynamics is a necessary condition for reliable EXC neuron firing.

Next, we asked whether this model could predict the results of our photostimulation experiments. We simulated optical activation by injecting a brief train of depolarizing current into SST units, with similar temporal properties as our experiments. As in our experiments, increasing the strength of SST inhibition increased EXC reliability and suppressed PV units (Fig. 5f, h). Notably, our model demonstrates that SST activation is most effective at increasing reliability when EXC units were unreliable. We found a strong correlation between the change in EXC reliability and the change in PV unit activity, such that large increases in reliability were accompanied with a strong suppression of PV unit activity (Fig. 5g). Changes in EXC unit firing rate, on the other hand, were poorly predictive of the increase in reliability (Supplementary Fig. 9). Removing the SST→PV circuit reduced the SST-induced suppression of PV units (Fig. 5h), and increased EXC variability (Fig. 5i). Thus, these simulations suggest that SST-INs increase reliability primarily by suppressing PV-INs.

Our model also correctly predicted a decrease in reliability following PV activation (Fig. 5j). Removing the SST→PV circuit did not affect these results (p = 0.12, Kruskal-Wallis one-way ANOVA). These results suggest that PV units might be injecting noise into EXC units. In support of this idea, transiently suppressing PV units by injecting a brief hyperpolarizing current increased reliability when EXC units were unreliable, but decreased reliability when EXC units were reliable (Fig. 5k). Therefore, our model supports the hypothesis that SST-INs reduce variability in EXC neurons by suppressing PV-INs through the SST→PV circuit.

### Suppressing PV-INs improves EXC neuron reliability

The main prediction of our model is that the increase in EXC reliability is caused by a SST-induced suppression of PV-INs. To examine this prediction *in vivo*, we directly suppressed PV-INs in mice that transgenically expressed Arch (PV-Cre x Ai35 mice, Fig. 6a). As expected, Arch-expressing PV-INs were strongly suppressed following green laser stimulation, with a latency that was comparable to SST activation (Fig. 6b). This method therefore mimicked the suppressive effect that SST activation had on PV-INs while avoiding the direct effect of SST inhibition on EXC neurons.

**Figure 6.**
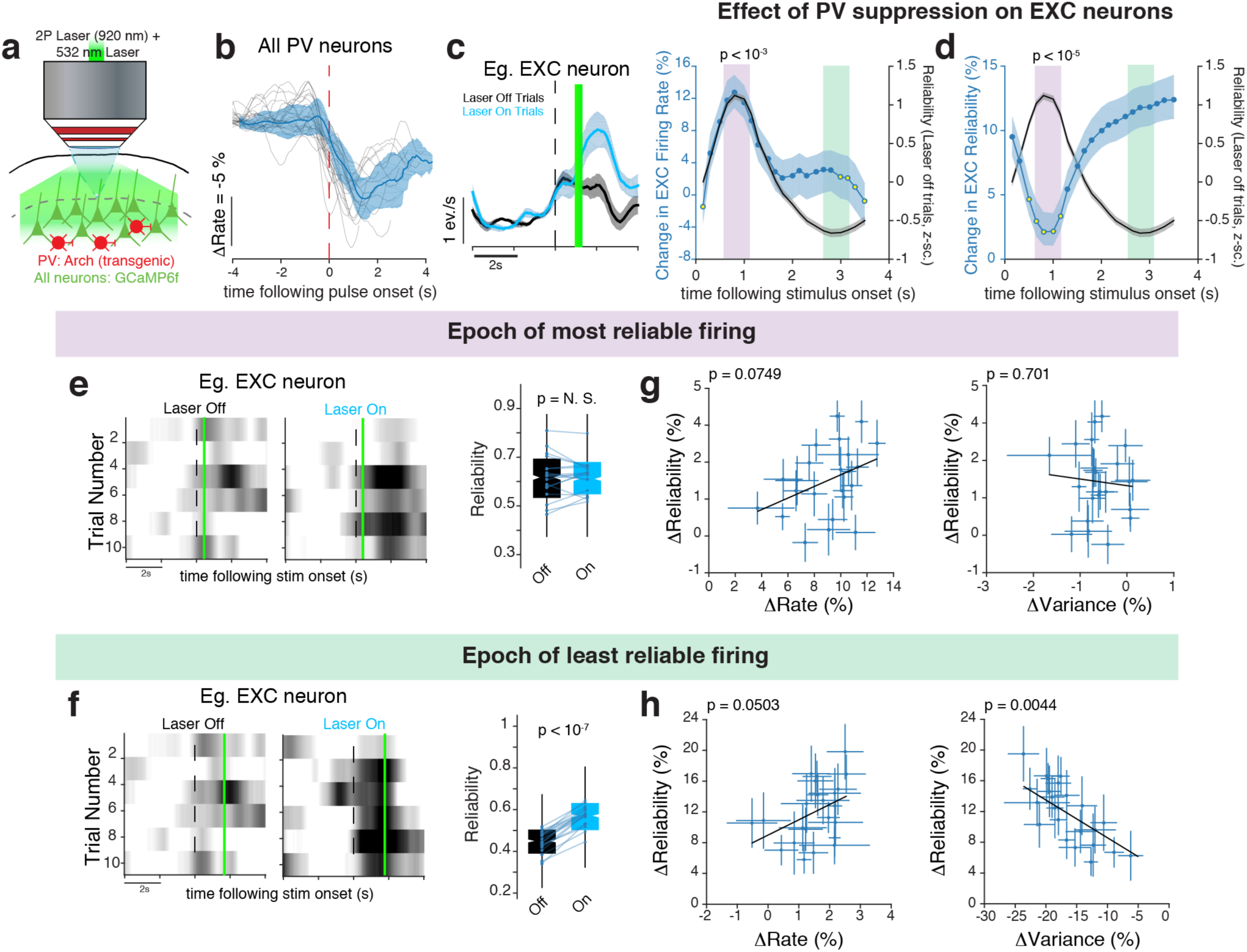
Suppressing PV-INs increases EXC neuron reliability. **(a)** Experimental setup. **(b)** Arch activation transiently suppresses PV-INs with short latency following laser onset. **(c)** Left: Suppressing PV-INs transiently increases the response rate of EXC neurons. Right: Change in firing rate for all PV suppression event. **(d)** Change in EXC neuron reliability, aligned to reliability on *Laser-off* trials. **(e)** Left: Representative raster plot of an EXC neuron showing no change in reliability following PV suppression during epoch of most reliable firing. Right: Box-whisker plot summarizing the effect of PV suppression. Each dot represents the median reliability from each imaged population. P-value computed using Bonferroni-corrected rank-sum test. **(f)** Same as **(e)**, but showing an increase in reliability following PV suppression during epoch of least reliable firing. **(g)** Changes in reliability that occur when PV-INs are suppressed during epoch of most reliable firing are weakly due to ΔRate (left) but not ΔVariance (right). Each data point in I and J shows median change for each imaged population. Error-bars are the 95% CI of the median. P-values computed from multivariate linear regression analysis. **(h)** Same as **(g)**, but shows that the increase in reliability is strongly associated with a reduction in variance but not a change in rate. All data in this figure are from 8 mice (634 neurons, 22 populations).

Due to a transient lifting of somatic inhibition, optically suppressing PV-INs strongly increased response rates when EXC neurons were most active (Fig. 6c). Despite this increase in response rate, suppressing PV-INs during epochs of reliable firing did not significantly change either reliability or between-trial variance (Figs. 6d, e). On the other hand, reducing PV inhibition during epochs of unreliable firing increased EXC neuron reliability (Fig. 6f), similar to SST activation. This change in reliability was primarily due to a reduction in between-trial variance as ΔRate was not a statistically significant predictor (Fig. 6g, h). Under control conditions, the green laser alone was unable to change either the firing rate or the reliability of EXC neurons (Supplementary Fig. 6). Therefore, transiently reducing PV inhibition with Arch had a similar effect on EXC neuron reliability as increased SST-IN activity, confirming our prediction.

Collectively, these experimental and computational results support a model, summarized in Supplementary Figure 10, in which PV and SST-INs work together to modulate the reliability of EXC neurons. In particular, an increase in SST activity and a corresponding suppression of PV-INs during can increase reliability, and an increase in PV activity can decrease reliability.

### Increasing reliability improves the ability to discriminate between uncertain stimuli

It has been noted that trial-to-trial fluctuations in behavioral performance correlate with neuronal variability in higher cortical areas^41^. However, the impact of unreliable coding, especially in V1, on perception remains unknown. Specifically, we asked whether modulating the reliability of V1 neurons could directly influence visual perception. To answer this question, we trained PV-ChR2 and SST-ChR2 mice on a Go/No-Go natural movie discrimination task. Specifically, water-restricted mice learned to discriminate between a target and a non-target movie (each 2s long) to gain a water reward (Fig. 7a). Wrong choices were punished with a brief acoustic white noise burst. We assessed performance by quantifying the number of correct responses, which is the total number of hits (licks to a target movie) and correct rejects (licks withheld to the non-target movie). Mice were trained progressively until they became proficient at associating the target movie with a water reward, and were typically able to perform this task within 18-21 days (10 mice). Post-training, mice were able to maintain a high performance (Hit > 70% and Correct Reject > 70%) over several sessions (400-500 trials/session, Supplementary Fig. 11). Mice that failed to meet this criterion were excluded from further analysis.

**Figure 7.**
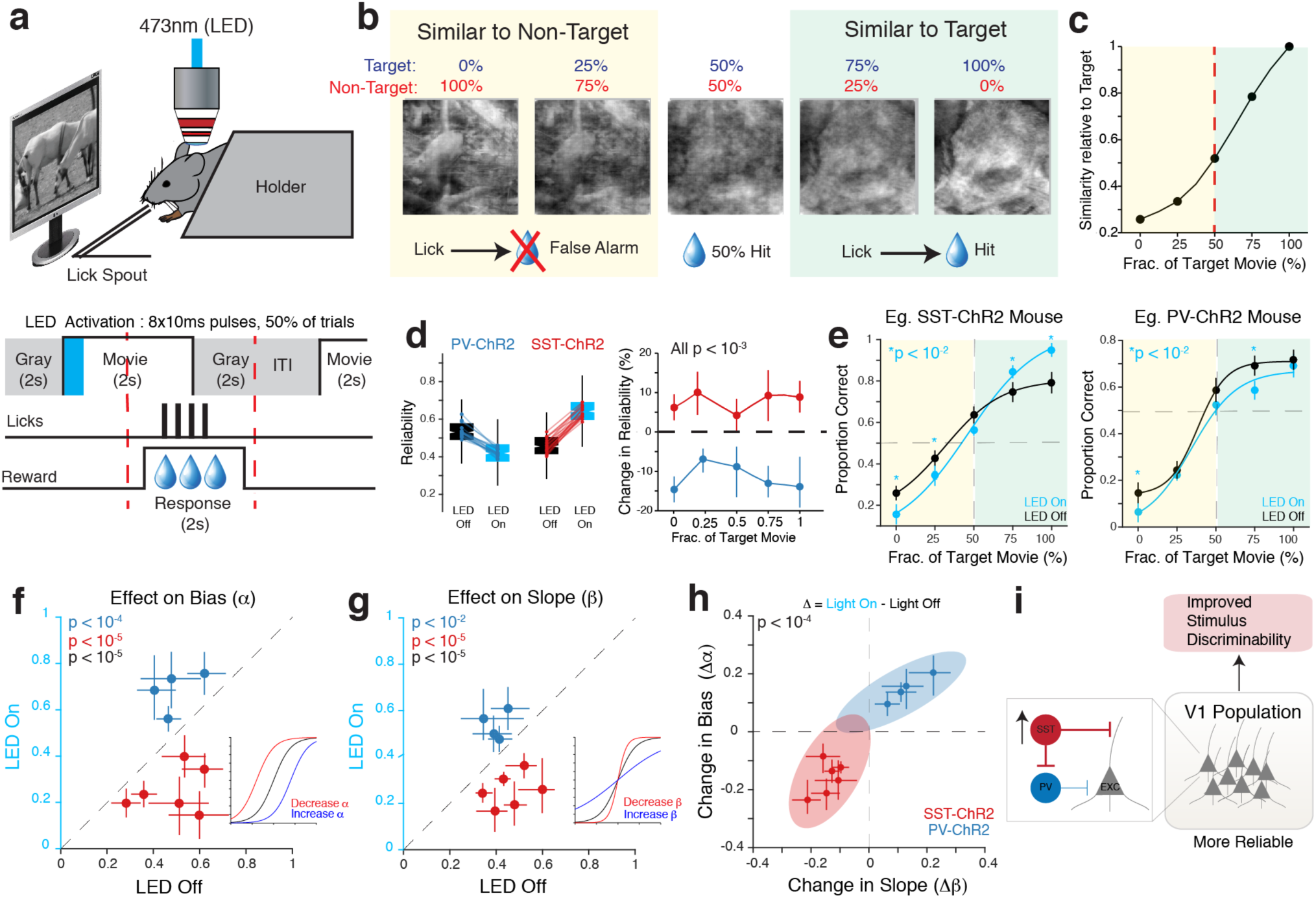
Activating SST-INs improves the ability of mice to discriminate complex stimuli. **(a)** Top: Experimental set up. Bottom: Schematic showing the timing structure of one trial. Licking during the response period window following a target movie resulted in a water reward. Licking outside the response window, or following a non-target movie, was not rewarded. **(b)** Example frames from phase blended movies. (c) Similarity between these movies and the target movie was quantified using the structural similarity index. Increasing the fraction of the target movie in the blend increased the similarity index monotonically. Non-target-biased movies (yellow shaded) were less similar, while target-based movies (green shaded) were more similar to the original target movie. (d) Left: Box-whisker plots showing a change in EXC neuron reliability for all movies following PV and SST activation in untrained mice. Each dot represents the median reliability of an imaged population of neurons. In both conditions, p < 10^−4^ (Bonferroni-corrected rank-sum test) compared the *LED-off* condition. Right: SST activation improves, and PV activation decreases EXC reliability for each phase-blended movie. All changes in reliability are significant (p < 10^−3^, Fisher’s Exact Test for each movie relative to 0). Data from, PV-ChR2: 3 mice (98 neurons) and SST-ChR2: 3 mice (121 neurons). **(e)** Psychometric curves for two mice comparing the effect of SST and PV-IN activation on the proportion of correct responses to the different phase-blended movies. The solid lines show the logistic function that best fit the data points. Data points are the average proportion correct and the error bars are the SEM over all sessions. Left: Activating SST-INs increased lick probability to the target movie, and decreased licking to the non-target movies, resulting in a sharper psychometric function (blue line). Right: Activating PV-INs decreased licking for most stimuli, resulting in a flatter psychometric function. P values (indicated in figure) computed using oneway Kruskal-Wallis ANOVA between *LED-on* and *LED-off*. **(f-g)** Scatter plots comparing the effect of activating PV (blue) and SST-INs (red) on the bias **(f)** and slope **(g)** of the psychometric function. Insets illustrate the effect of changing the bias and slope in a toy psychometric function. Each dot represents data from a single mouse and error-bars denote SEM over all sessions for that mouse. P-values computed using Friedman’s test (effect of laser vs. effect of IN activation) followed by post-hoc tests. The comparisons tested are: *LED-on* vs. *LED-off* (blue, PV-ChR2 and red, SST-ChR2) and difference for PV-ChR2 vs. difference for SST-ChR2 (black). **(h)** Scatter plot comparing the change in bias and slope following PV and SST activation. Activating SST-INs sharpened the slope and decreased the bias, whereas activating PV-INs had the opposite effect. Shaded ellipses represents the 95% CI over all sessions. Differences between SST and PV activation are significant, p < 10^−4^ (Friedman’s Test). All data in from F-H are from 6 SST-ChR2 mice and 4 PV-ChR2 mice. **(i)** Summary.

Once proficient, we increased the complexity of the task by altering the spatial statistics of both movies. It is well established that the phase spectrum of natural scenes is essential in image recognition because it contains salient structural information, such as the location of edges ^42^. We used this fact to introduce uncertainty in the target movie by blending its phase spectrum with the phase spectrum of the non-target movie at different fractions (Fig. 7b). We also equalized the amplitude spectra of these surrogate movies to the mean spectrum of the target and non-target movies (Supplementary Fig. 12a). All other image statistics (contrast, luminance, etc.) were also fixed between movies. As a result, mice could only use subtle differences in phase information to discriminate between these movies. Importantly, we observed no difference in EXC neuron reliability in naïve, untrained mice, between these movies (Supplementary Fig. 12b). Thus, in essence, mice had to perform a categorization task, where they assessed the similarity of the movie presented to the learned target movie (Fig. 7c). We reasoned that if increasing response reliability does improve stimulus selectivity, then mice should be able to correctly identify the rewarded target movie from “noisy” versions.

Expectedly, mice licked more to movies that were similar to the target and less to movies that were similar to the non-target movie (Fig. 7e). We fit each psychometric curve with a two parameter logistic function ^43^, and assessed changes in bias (α) and slope (1/β) following IN activation. By this convention, a rightward shift of the psychometric curve corresponds to an increase in α, whereas an increase in slope would correspond to a decrease in β (see insets in Fig. 7g, h). To determine the effect that IN activation had on the ability of mice to accurately categorize these movies, we pulsed a blue (470 nm) LED at movie onset, on 50% of trials (randomly selected, Fig. 7a). In naïve, untrained mice, activating PV-INs under the same conditions suppressed reliability, whereas SST activation increased reliability, for all phase-blended movies (Fig. 7d), consistent with our earlier findings. Based on these results, we hypothesized that increasing SST inhibition should be increase discriminability between the phase-blended stimuli, which in turn would sharpen the psychometric curve (decreased β). In contrast, the increase in unreliability following PV activation would decrease discriminability and increase false alarms, which would flatten the psychometric curve (increased β).

In agreement with this hypothesis, activating SST-INs sharpened psychometric curves (Fig. 7e), by decreasing bias and increasing the slope (Fig. 7f-h). These changes indicate that increasing SST inhibition lowered the detection threshold and increased movie discriminability. Notably, these changes were due to an increase in hit rate and a decrease in false alarm rate (Supplementary Fig. 12c-e), suggesting that mice made fewer mistakes following SST activation. In contrast, activating PV-INs flattened the psychometric curve (Fig. 7e) by increasing bias and decreasing the slope. Increasing PV inhibition also significantly decreased hit rates without changing false alarm rates, indicating that the mice were less able to distinguish the target from the non-target movies.

Taken together, our results show that, by reducing neuronal variability within V1, SST-IN activation improves the ability of mice to recognize “noisy” versions of the target movie, whereas increasing PV inhibition is detrimental to performance (summarized in Fig. 7i).

## DISCUSSION

Reducing trial-to-trial variability within cortical neuron networks is critical for accurate sensory information processing; however, the underlying neural mechanisms remain unknown. In this study, we used novel double transgenic mice and all-optical physiology to reveal a previously unknown role of the SST→PV circuit in bi-directionally modulating the reliability EXC neurons to naturalistic stimuli in mouse V1.

Our experiments reveal that a necessary condition for reliable sensory processing is active SST-INs and weaker/suppressed PV-INs, and that this mutual antagonism is maintained through the inhibitory action of the SST→PV circuit. Surprisingly, a recent study identified PV-INs, but not SST-INs, as critical regulators of reliability. A key reason for this difference is that Zhu and colleagues suppressed PV-INs for a much longer duration than our study (6 s vs. 110 ms). The main advantage of our photo-stimulation method is that it allowed us to show that the effect of SST and PV-INs on modulating EXC variability is highly dependent on the current reliability of EXC neurons. Namely, SST-INs were less effective at increasing reliability during epochs of reliable firing, while PV-INs were less effective at reducing reliability during epochs of unreliable firing. Furthermore, our model simulations showed that without the SST→PV circuit, SST-IN activation decreased reliability. Our data therefore supports the idea that SST and PV-INs must provide temporally restricted inhibition in relation to EXC neurons to change variability. Long term suppression of PV and SST-INs as would likely disrupt this relationship between these interneurons. Therefore, our findings, together with others^27,44,45^, underscores the importance of using precisely timed perturbations to study the dynamics of cortical inhibition. Importantly, we demonstrate that the responsibility of modulating response reliability does not lie exclusively with one IN subtype; instead, it is the co-operative dynamics between SST and PV-INs, which is important for controlling the temporal fidelity of sensory processing.

Previous work has shown that feed-forward inhibition, acting through fast-spiking PV-INs, plays a critical role in shaping the temporal fidelity of EXC neurons. For example, the delay between inhibition and excitation creates a temporal integration window^14,22^ and variations in the duration of this window changes the spiking precision of EXC neurons to sensory stimulation^46^. However, sparse activity patterns, which are common during natural scene stimulation^47^, strongly recruit recurrent inhibition from SST-INs and only weakly recruit PV-INs^48^. Our work reconciles these observations and demonstrates how the SST→PV circuit allows recurrent inhibition to modulate the strength of feed-forward inhibition during epochs of reliable coding under naturalistic conditions.

We propose that a potential biophysical function of the SST→PV circuit is to maximize the signal-to-noise ratio of EXC neurons by minimizing noise in the synaptic inputs and maximizing spiking at the soma. Specifically, SST-INs are ideally poised to alter synaptic integration in EXC neurons by altering the active properties of dendrites in a branch-specific manner ^23,24,49,50^. This, in turn, would allow only the most reliable inputs to be integrated^16,51^. Our observation that SST-INs lag behind PV-INs during periods of reliable firing implies that, during these epochs, inhibition is routed away from the soma and into the dendrites. Similar results have been observed in the hippocampus and the barrel cortex, where inhibitory inputs shift from the soma to the dendrite depending on the firing rate of the neuron^48,52^. Computational models have shown that this mechanism allows SST-INs to adaptively adjust the integration threshold at the soma, which in turn can increase the robustness of spiking in the presence of stochastic inputs^53,54^. Future studies should be aimed at using our dual labeling technique to further characterize the interactions between PV-SST INs during timescales more relevant to synaptic integration. There is also a growing body of evidence that basal forebrain cholinergic inputs^55^, long-range excitatory inputs from other cortical areas^56,57^ can modulate SST-IN activity. Therefore, the SST→PV circuit is an appropriate target for top-down factors, such as arousal and attention, to alter local computations in V1 by changing EXC neuron variability.

The impact that sensory processing variability has on visual perception remains highly debated. Although several studies have established a relationship between trial-to-trial fluctuations in sensory neurons and perceptual decisions^41,58^, however, it is still unclear if these fluctuations can be filtered out at later processing stages^59^ and how this affects perception. Our behavioral results reveal that reducing variability, both in individual neurons and across the network, improves the discriminability of complex stimuli. In particular, we observed a marked decrease in false alarm rates following SST-IN activation, suggesting that mice were making fewer incorrect choices when neural reliability increased. Also, by manipulating the statistics of the target movie, we were able to show that SST-IN activation allowed mice to more effectively detect perturbations in stimulus appearance. Interestingly, a previous study found that activating PV-INs for long durations improved orientation discrimination in mice ^60^, and attributed this to an increase in both orientation selectivity and signal-to-noise ratio in EXC neurons. A key reason for this difference could be the change in circuit dynamics caused by long-duration PV stimulation^61^. However, despite this discrepancy in mechanism, both studies point toward a common theme that improved coding fidelity in V1, either through increased reliability or sharper selectivity, or both, can improve visual perception. This notion is further bolstered by several recent findings that both cholinergic modulation and higher cortical feedback, which also change response reliability and selectivity, can similarly improve stimulus discriminability^56,62^. Furthermore, through our novel categorization task, we are able to demonstrate, for the first time in mice, that reducing response variability early in sensory processing can also decrease perceptual uncertainty, as previously predicted by theoretical models^63^.

In conclusion, our study establishes that PV and SST-INs have complementary roles in controlling neuronal response reliability. The cooperative action of these INs provides a powerful computational mechanism by which response variability can be titrated based on task demands and internal state to improve the coding of stimulus information. In addition to the visual system, this strategy could also be active in other cortical areas to effectively gate the flow of information.

## SUPPLEMENTARY FIGURES AND LEGENDS

**Supplementary Figure 1.**
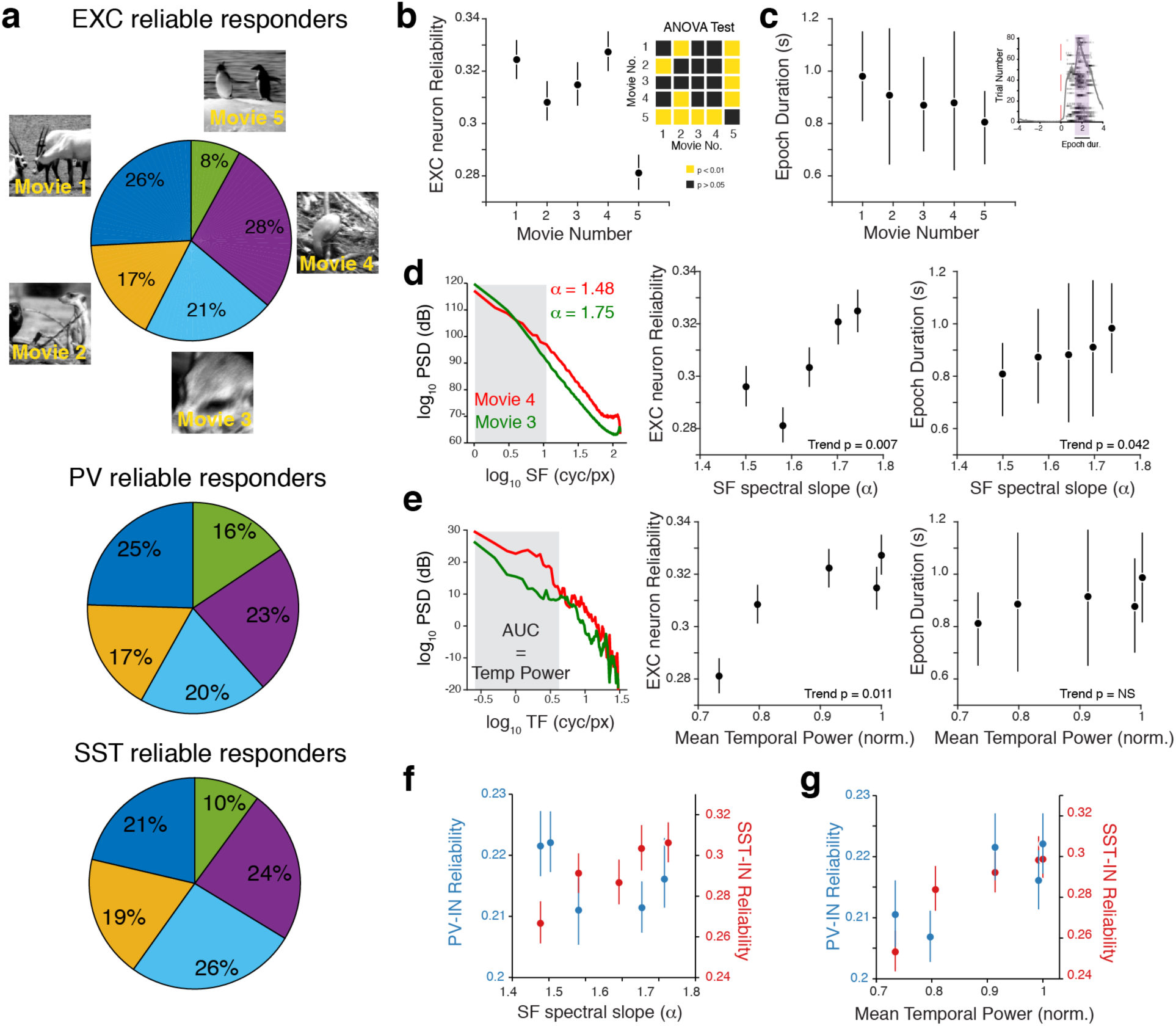
Reliability values differ from between movies and depend on spatio-temporal statistics. **(a)** Pie charts showing movie-wise distribution of reliably responding EXC (Top), PV (Middle) and SST (Bottom) neurons. Movies which recruit a greater fraction of reliably responding EXC neurons also recruit more reliably responding PV and SST-INs. **(b)** Movie-wise distribution of EXC neuron reliability. Inset, one-way ANOVA table showing pairs of movies with significantly different reliability values. **(c)** Movie-wise distribution of the duration of reliable epoch. The epoch duration (see inset) was defined as the length of time with reliability >0.4 for each neuron. **(d)** The spectral slope, which is a measure of spatial correlations between pixels was calculated from the spatial power spectral density of each movie as described previously^33^. Movies with higher spectral slope (stronger spatial correlations) had larger reliability values and longer epochs of reliable spiking. **(e)** The temporal power, which is a measure of the strength of temporal correlations between pixels, was computed by integrating the temporal power spectral density (left). Movies with stronger temporal correlations also had larger reliability values. **(f)** SST-INs respond more reliably in movies that have higher spectral slope. **(g)** Both PV and SST-INs respond more reliably in movies that have higher temporal correlations. All data represented as mean +/- SEM. Data from: PV = 8 mice (690 neurons); SST = 8 mice (368 neurons); EXC = 10 mice (1101 neurons). All trend P values computed using linear regression (F-test relative to constant model).

**Supplementary Figure 2.**
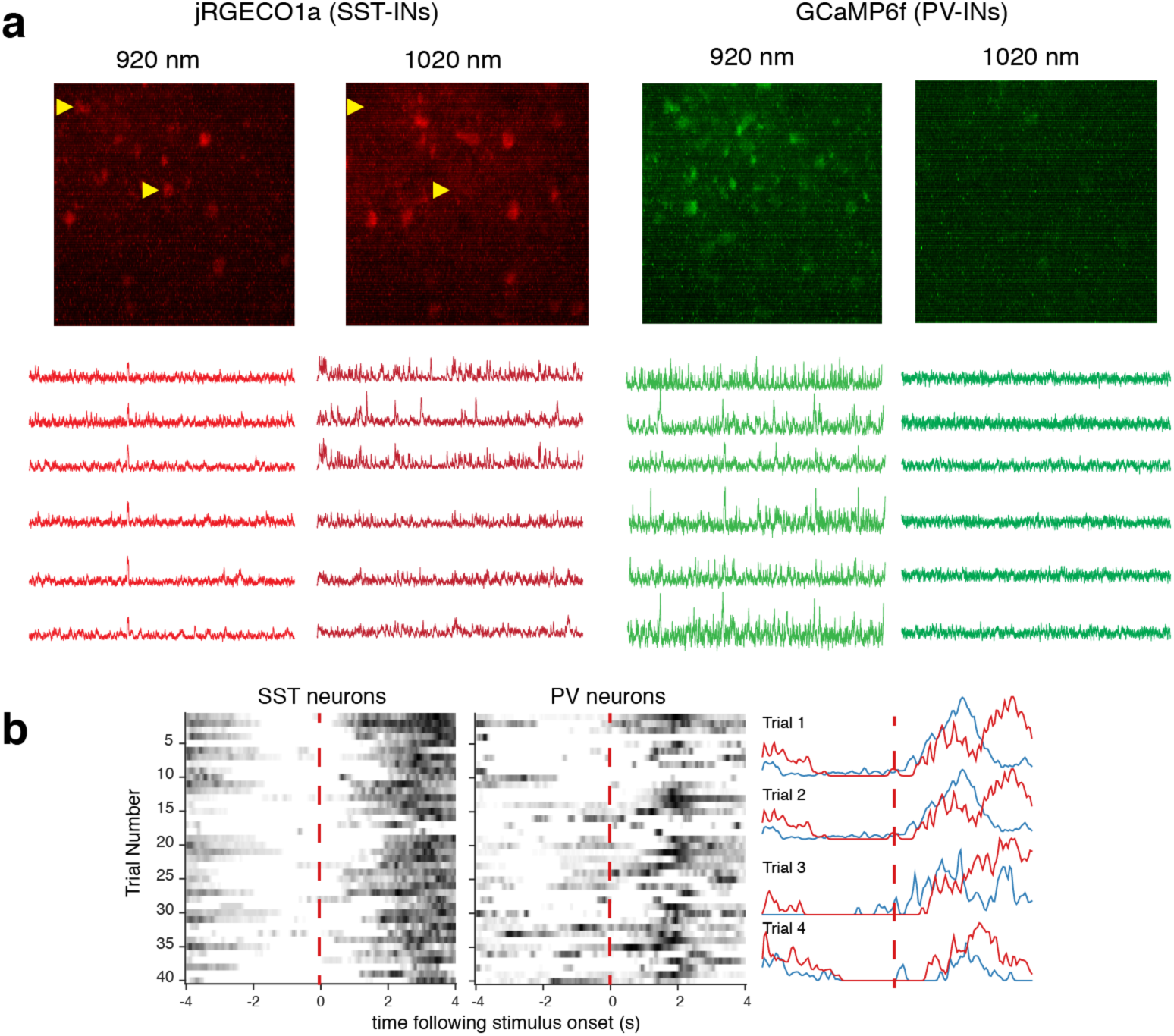
Dual wavelength imaging of PV and SST neurons. **(a)** *Top:* Images of jRGECO1a-expressing SST-INs and GCaMP6f-expressing PV-INs taken at 920 nm and 1020 nm respectively. Bleed through from the green to the red channel can be clearly seen at 920 nm (indicated by yellow arrowheads). In contrast, no green signal can be detected at 1020nm. *Bottom:* No jRGECO1a activity can be detected at 920 nm compared to 1020 nm. In contrast, no GCaMP6f activity can be detected at 1020 nm. Each trace is matched to the same neuron and shows activity in response to a series of natural movies (800 s long, acquired at 20 Hz). **(b)** Example showing trial-by-trial correlation between PV and SST-INs.

**Supplementary Figure 3.**
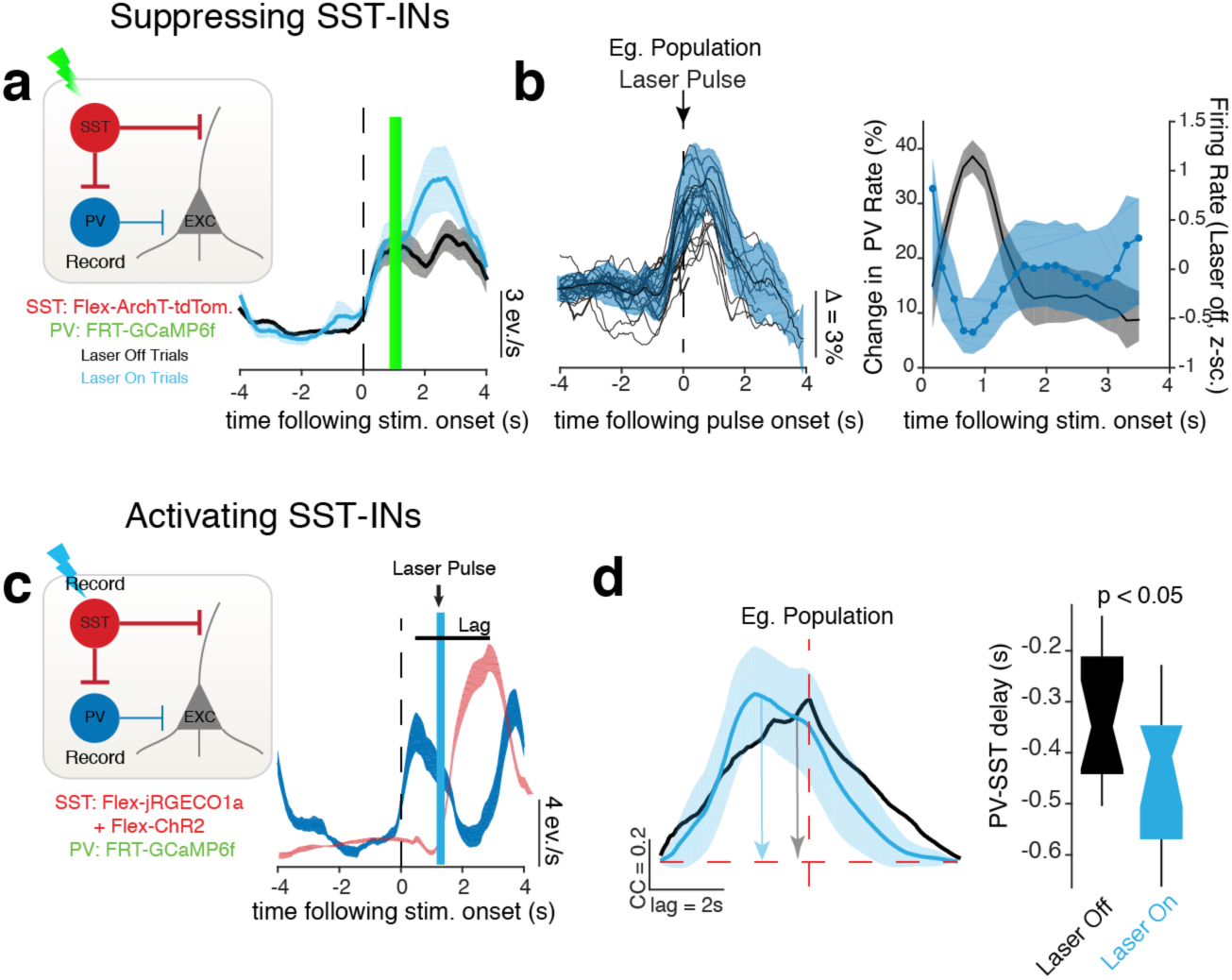
SST-INs strongly inhibit PV-INs via the SST→PV circuit. **(a)** Inset, Cre-dependent ArchT was expressed in SST-INs while Flp-dependent GCaMP6f was expressed in PV-INs in SXP mice. Representative example trial-averaged firing rate from one PV-IN showing that suppressing SST-INs strongly increases the rate of PV-INs. Shaded area, SEM over trials. **(b)** Left: Response rate change in one representative population of PV-INs (8 neurons) aligned to laser onset. All PV-INs increase their firing rate following suppression of SST-INs. Right: Quantification of change in response rate of PV neurons following SST suppression relative to response rate on laser-off trials. We observe a significant increase in PV activity (p < 0.001, permutation test) regardless when SST-INs are suppressed during a movie‥ Shaded area, 95% CI. Data from 3 mice (121 PV neurons). **(c)** Inset, Cre-dependent ChR2 and jRGECO1a was expressed in SST-INs while Flp-dependent GCaMP6f was expressed in PV-INs in SXP mice. Representative example trial-averaged firing rate from one simultaneously imaged PV-SST pair, showing a strong suppression of PV-INs following SST-IN activation. The peak suppression occurs almost at the same time as SST-INs reach peak activation. **(d)** Left: Example cross-correlogram between all pairs of simultaneously recorded SST (n = 4) and PV-INs (n = 9) for an example population, showing the effect of SST activation on the time lag between PV and SST-INs. Gaussian fit is not shown. Data here is averaged over all stimulation epochs. Shaded area, SEM. Right: Box-whisker plot showing that activating SST-INs increases the time lag between PV and SST-INs (p = 0.235, Bonferroni-corrected rank-sum test). Data from 3 mice (84 PV neurons, 39 SST neurons).

**Supplementary Figure 4.**
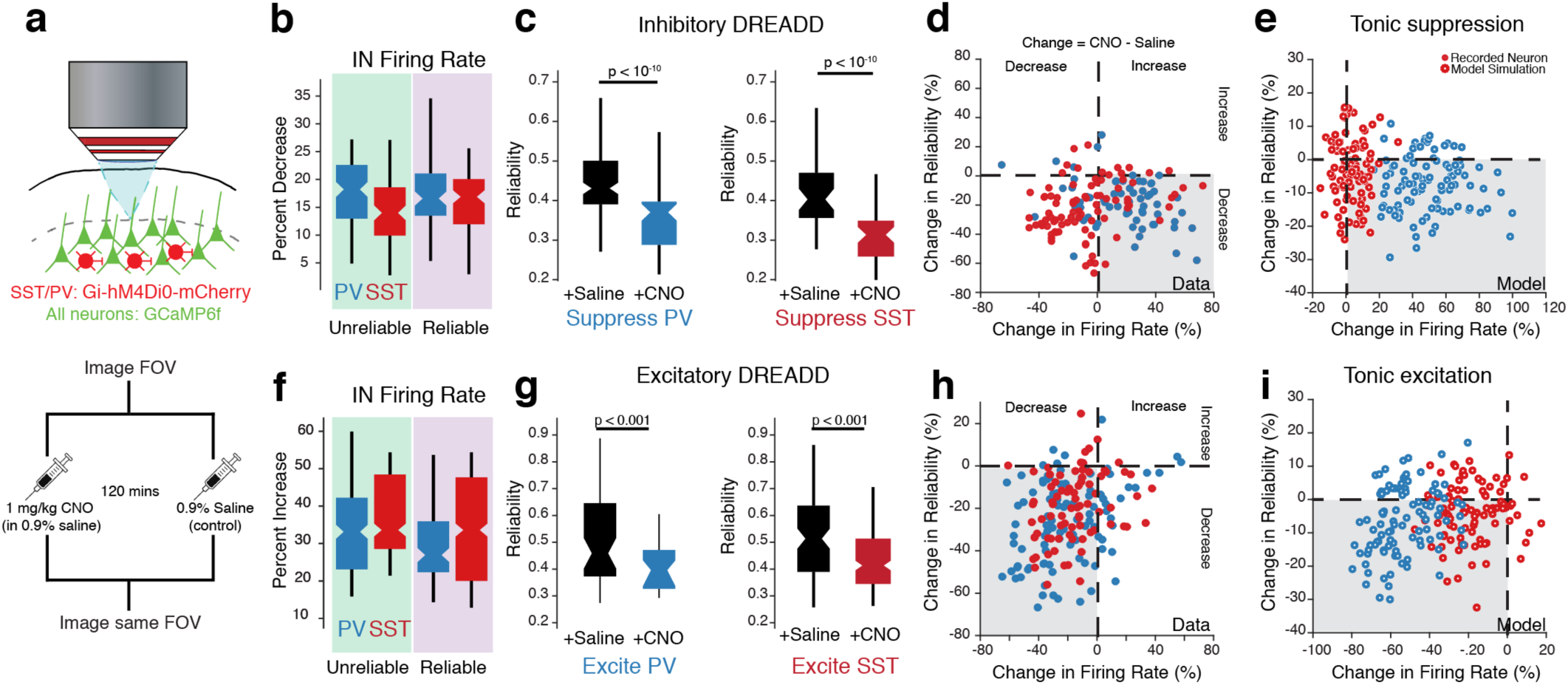
Chronic, chemogenetic perturbation of both PV and SST-INs reduces EXC neuron reliability. **(a)** Schematic describing experimental set up. **(b)** Percent firing rate decrease in PV and SST-INs expressing inhibitory DREADD following CNO injection (compared with saline controls). As with Fig. 1f, separately analyzed changes in firing rate during epochs of unreliable or reliable EXC neuron firing (EXC and INs (mCherry expressing) were imaged simultaneously). This analysis shows that inhibitory (Gi) DREADD non-specifically reduced the firing rates of PV and SST-INs. **(c)** Box-whisker plots showing the effect of suppressing PV and SST-INs on EXC neuron reliability. **(d)** Scatter plot relating the change in reliability to the DREADD-induced change in firing rate. The change in reliability was not due to a change in firing rate as neuron that either increased or decreased their firing rate had reduced reliability following CNO administration. **(e)** Model simulation with tonic IN suppression can reproduce the DREADD-induced decrease in reliability. **(f-i)** Same as (b-e) but with Excitatory (Gq) DREADD instead. Data from: 3 mice each (SST-Gi-DREADD: 20 SST, 64 EXC; SST-Gq-DREADD: 23 ST, 52 EXC; PV-Gi-DREADD: 38 PV, 72 EXC; PV-Gq-DREADD: 16 PV, 89 EXC). All P-values computed using one-way Kruskal-Wallis ANOVA.

**Supplementary Figure 5.**
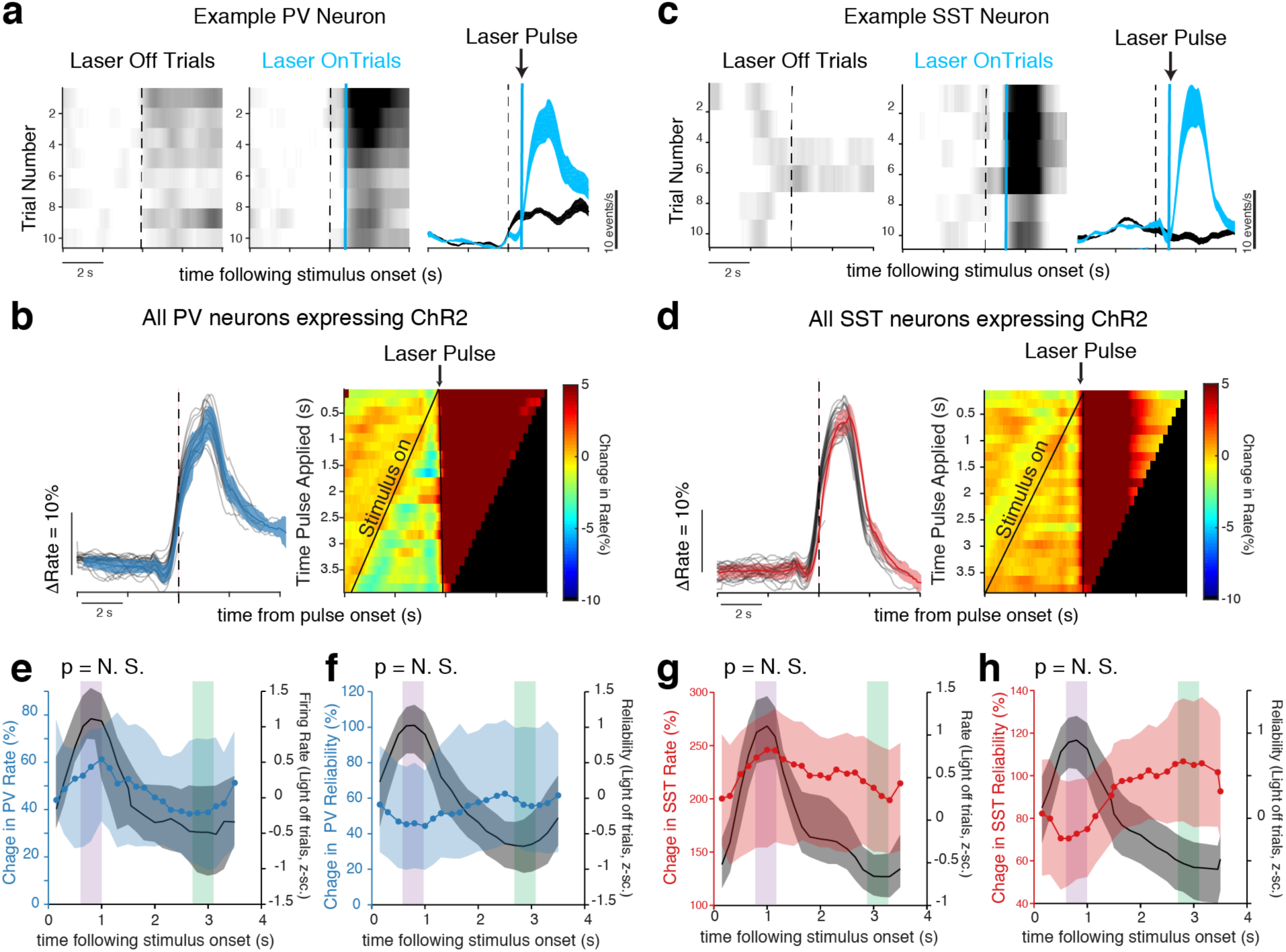
Laser activation reliably activates both PV and SST-INs. **(a-b)** Example raster plot showing the effect of laser activation on a PV-IN expressing both GCaMP6f and ChR2. Shaded area, SEM over trials. **(b)** *Left:* Response rate trace of the same PV-IN as in **(a)**, but aligned to laser onset instead. *Right:* Heat map showing the average change in response rate of all PV neurons relative to the *Laser-off* condition (n = 3 mice, 58 neurons), aligned to pulse onset. All laser pulses, regardless of when they are applied during the movie, result in a strong increase in firing rate. **(c-d)** Same as **(a-b)** but for SST-INs instead. These plots show that both PV and SST neurons are reliably and strongly activated following laser activation, regardless of when they are applied during the movie. **(e-h)** Quantification of firing rate **(e, g)** and reliability **(f, h)** change for all PV-and SST-INs. Data from: PV-Cre = 3 mice (58 neurons), SST-Cre = 3 mice (52 neurons). Shaded area represents 95% CI. All changes are significant p < 0.0001 (permutation test).

**Supplementary Figure 6.**
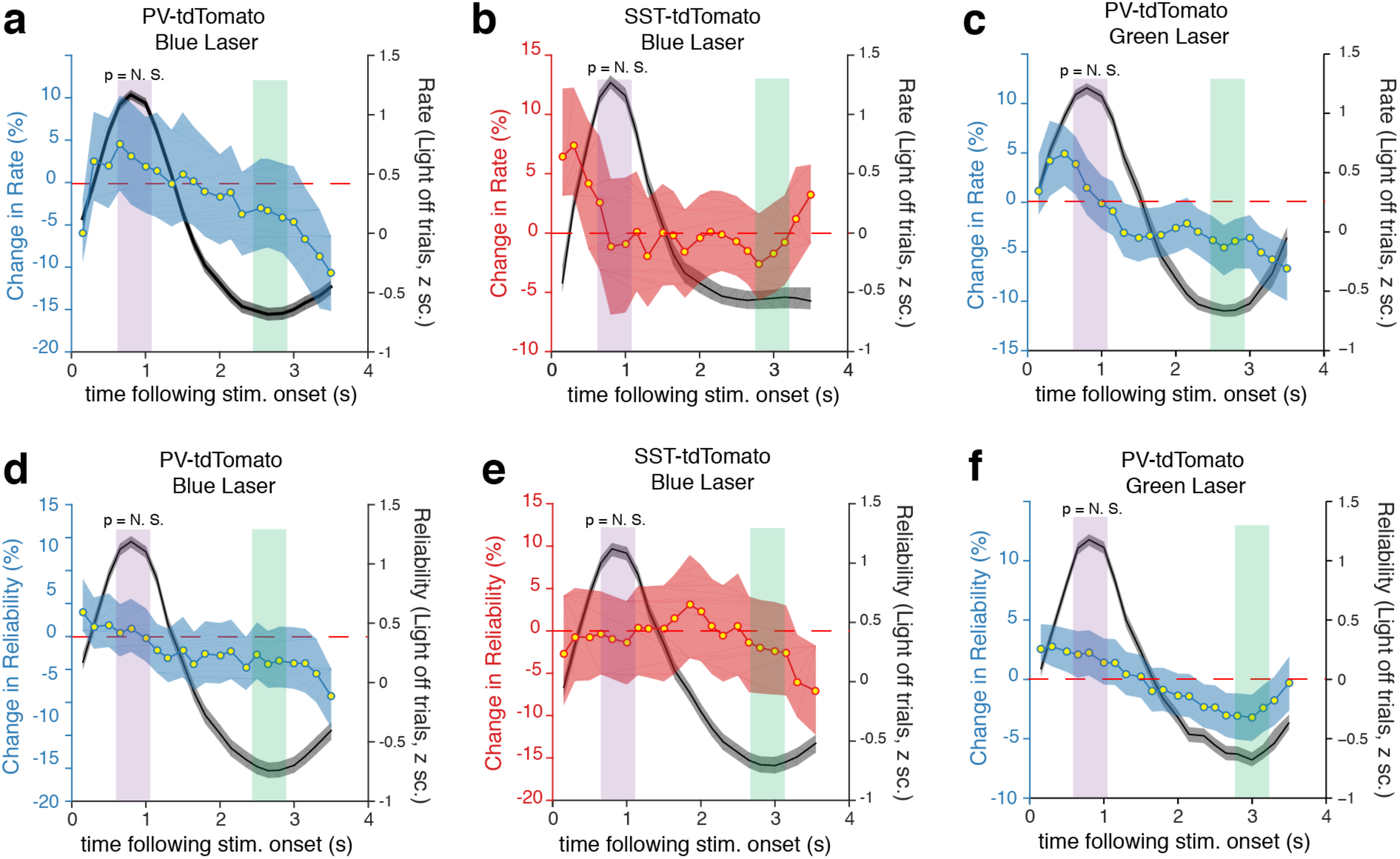
Change in rate and reliability is not due to stimulation laser artifacts. **(a-c)** No significant change in response rate in tdTomato-expressing mice following stimulation with either blue **(a, b)** or green laser **(c)**. For details, see Methods. **(d-f)** No significant change in reliability in the same tdTomato-expressing mice. Data pooled from: PV-tdTomato (blue laser) = 3 mice (120 neurons), PV-tdTomato (green laser) = 3 mice (102 neurons), SST-tdTomato = 3 mice (98 neurons). All p-values are non-significant (permutation test). Shaded areas represent 95% CI of median.

**Supplementary Figure 7.**
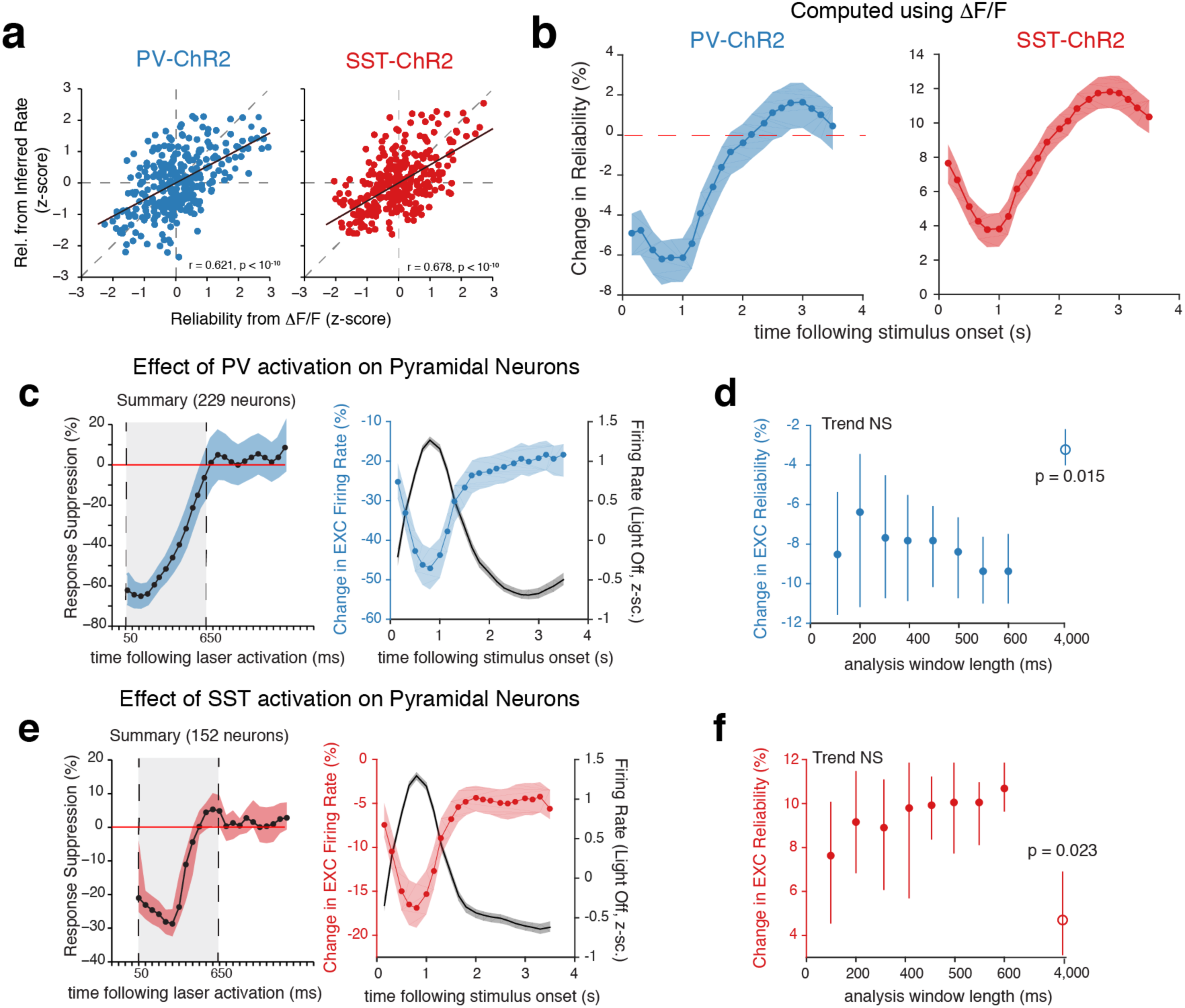
Deconvolution and analysis window length does not affect result. **(a)** Reliability computed from inferred firing rates is similar to reliability measured from raw fluorescence changes (DF/F). Scatter plot shows that neurons with reliable DF/F will also have reliable inferred firing rates. **(b)** Change in reliability measured using DF/F without deconvolution. Data same as Figs 3 and 4. **(c)** Percent change in EXC neuron response following laser activation of PV-INs. All data analysis was limited to a 600 ms window indicated by the gray box. During this period, the laser maximally suppresses pyramidal neurons. **(d)** Plot of change in EXC reliability following PV activation at stimulus onset over different analysis window lengths. We found that changing the window length within 50 (1 frame) to 600 (12 frames) ms following laser offset did not significantly affect the reduction in EXC reliability caused by PV activation. The effect, however was significantly diminished when the entire 4 s stimulus-on period (open circle) was included in the analysis. This is mainly due to the fact that PV-INs stop exerting their inhibitory effect on EXC neurons after 650 ms as shown in (c). **(e-f)** Same as **(c-d)** but for SST-IN activation instead. Again changing the duration of the analysis window does not affect the increase in EXC reliability caused by SST activation. Data in c-f shown as mean +/- SEM for stimulation at stimulus onset. Analysis of other stimulation epochs yielded qualitatively similar results. All shaded areas are 95% CI of median.

**Supplementary Figure 8.**
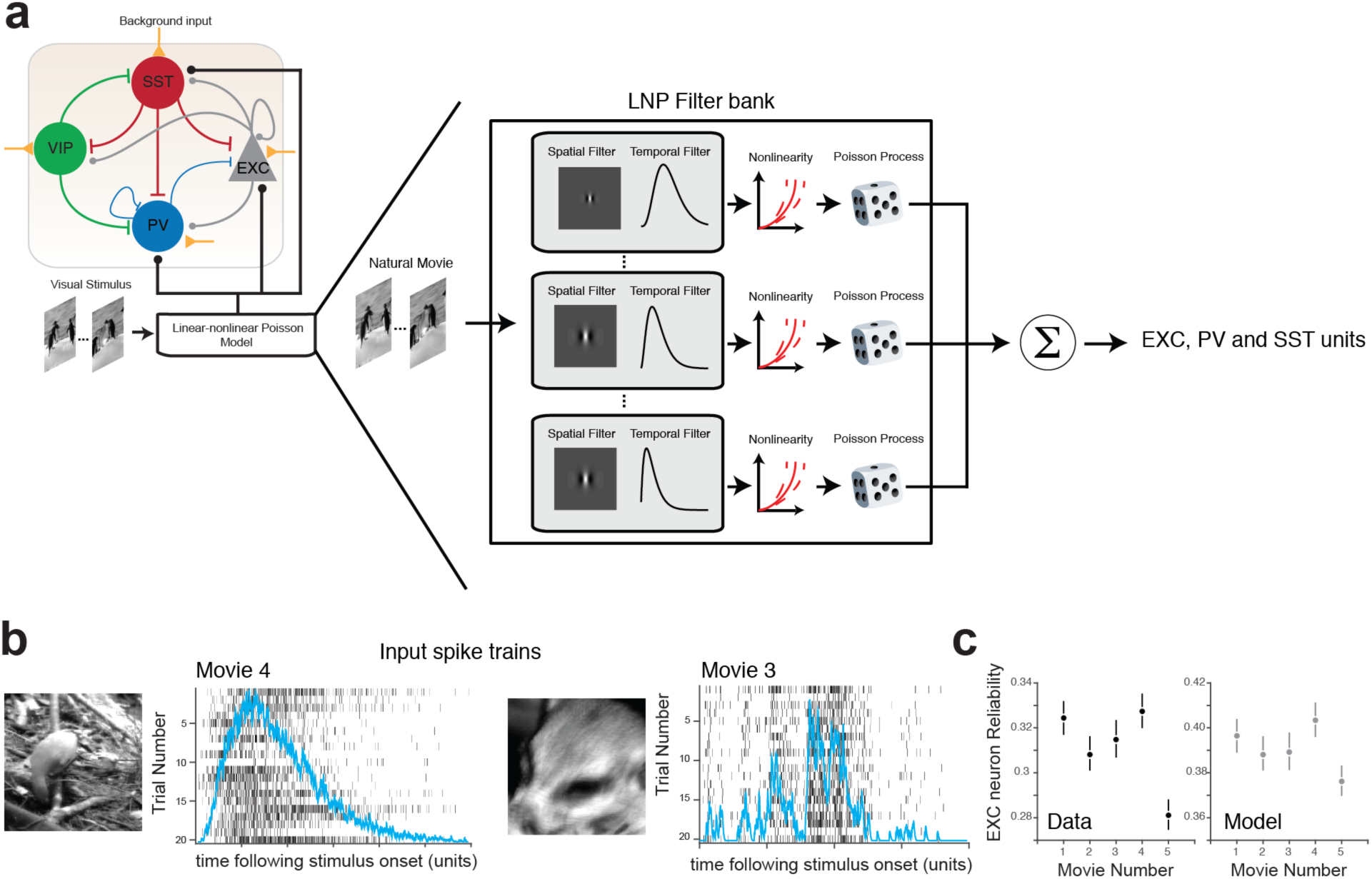
Linear-nonlinear Poisson cascade model to simulate visual input. **(a)** Schematic showing components of the linear-nonlinear-Poisson (LNP) model. Model is described in detail in the Methods section. **(b)** Example input spike trains produced by the LNP model along with their estimated firing rates (blue lines, normalized to maximum) to two different natural movies. Note that the model captures the different temporal properties of each movie, and, as a result, produces different inputs for each movie. For each movie, these spike trains are summed and used as an input to either EXC, PV and SST units. **(c)** The LNP model is able to recapitulate the movie-wise trend in EXC neuron reliability observed in the experimental data (black dots, same as Supplementary Fig. 1). The grey dots are the average reliability of EXC units in the model (from 500 simulations, see Methods). Errorbars, SEM.

**Supplementary Figure 9.**
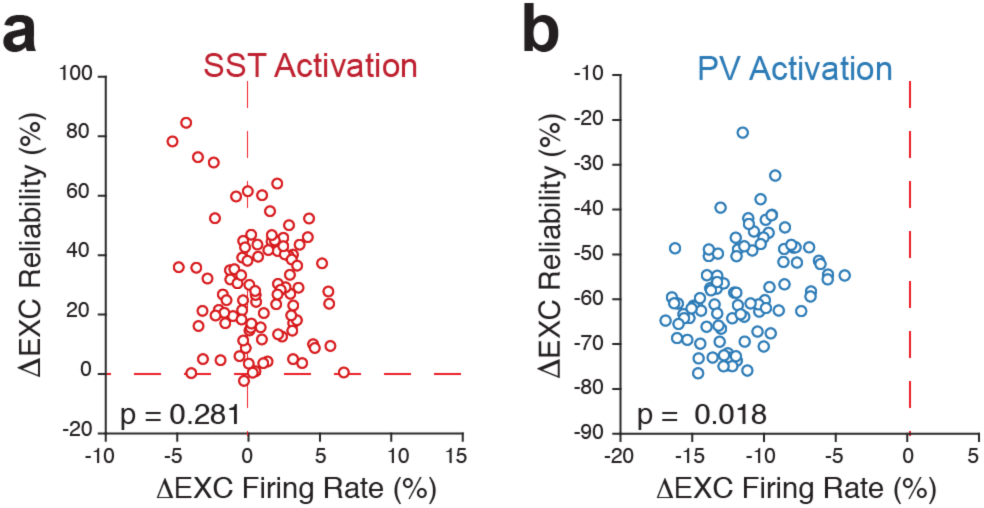
Changes in reliability due to SST activation are poorly predicted by changes in EXC firing rate. **(a)** Scatter plot showing no significant relationship between the change in EXC unit reliability and firing rate following SST unit activation. Each data point is an independent model simulation (see Fig. 5 and **Methods**). **(b)** In contrast, following PV unit activation, the change in EXC unit reliability can be predicted from a change in firing rate. P-values computed using multivariate linear regression.

**Supplementary Figure 10.**
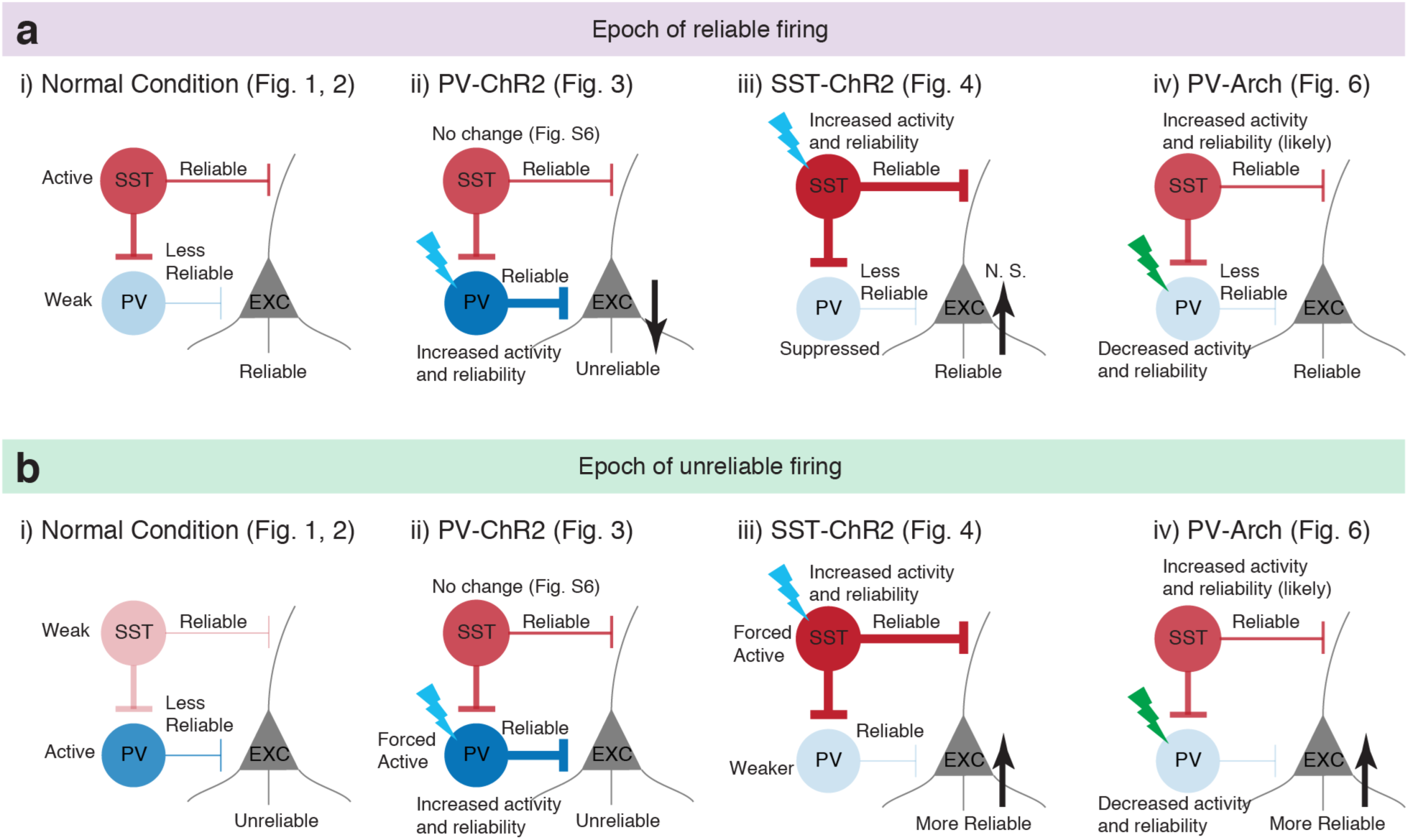
Cartoon summarizing main results of this paper. **(a)** Changes observed during epoch of reliable firing. **(i)** In the normal condition (i.e. with no external perturbations), SST neurons are more active and more reliable than PV neurons during periods of reliable EXC neuron firing. **(ii)** Activating PV neurons increases the activity and reliability of PV neurons, but does not change the activity of SST neurons (Supplementary Fig. 3). This leads to a decrease in EXC neuron reliability and an increase in variability across trials (Fig. 3). **(iii)** Activating SST neurons increases the activity and reliability of SST neurons, and decreases the activity and reliability of PV neurons (Fig. 4). This leads to marginal increase in reliability. **(iv)** Directly suppressing PV neurons strongly disinhibits EXC neurons but does not significantly alter reliability. Therefore, a hallmark of periods of reliable firing is increased SST-neuron activity relative to PV neurons because factors that increase SST activity increase reliability. **(b)** Same as **(a)**, but for changes observed during epoch of unreliable firing. This shows that a hallmark of periods of unreliable firing is increased PV-IN activity relative to SST-INs, because factors that increase PV activity decrease reliability, whereas factors that decrease PV activity increase reliability respectively.

**Supplementary Figure 11.**
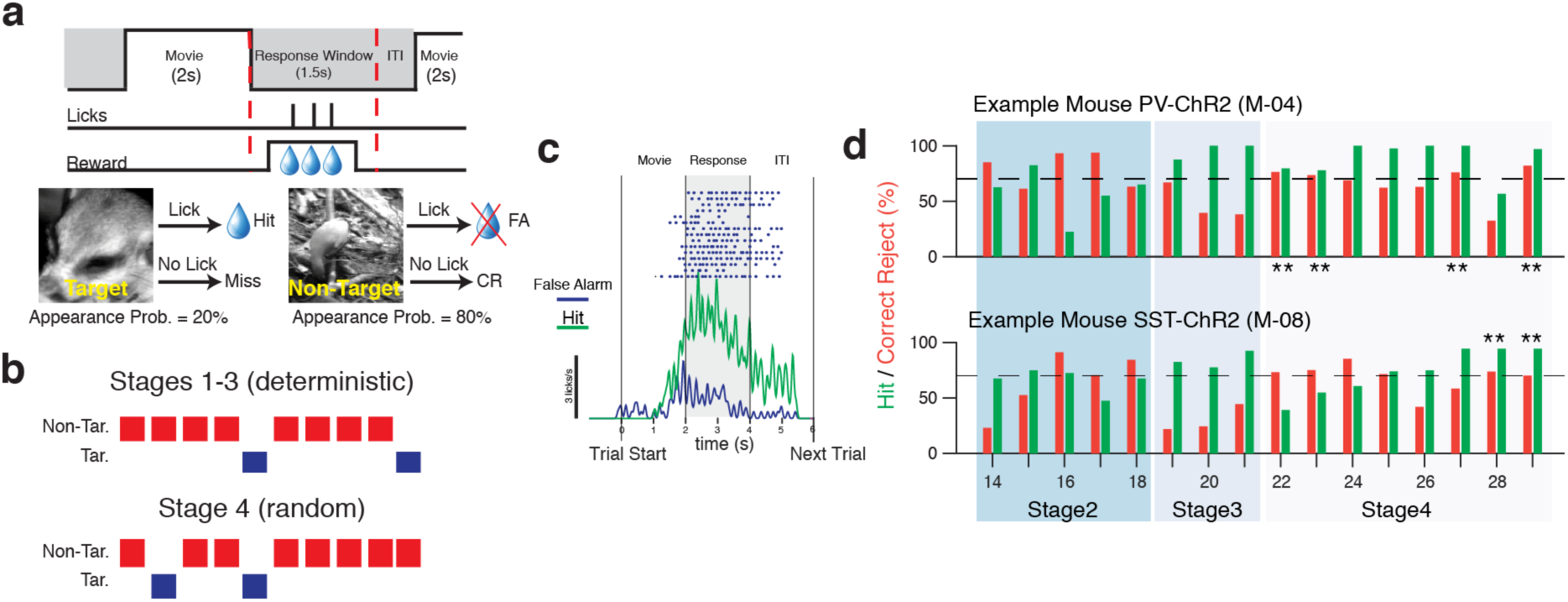
Natural movie discrimination task training. **(a)** Cartoon showing timing of stimuli and response window during training phase. **(b)** Cartoon illustrating order of stimuli during the various training stages (described in Supplementary Table 2). **(c)** Representative lick raster of a well-trained mouse (during Stage 4). Each dot denotes a lick, and colored lines denoted trial-averaged lick rates. From this example, it can be seen that well-trained mice restrict their licking only during the response period. False alarms are generally caused by anticipatory licking. **(d)** Correct Reject and Hit Rate performance of two representative mice during the course of training from Stage 2 to Stage 4. Each pair of bars corresponds to a training session. Stars indicate sessions in which performance crossed the threshold (dashed line).

**Supplementary Figure 12.**
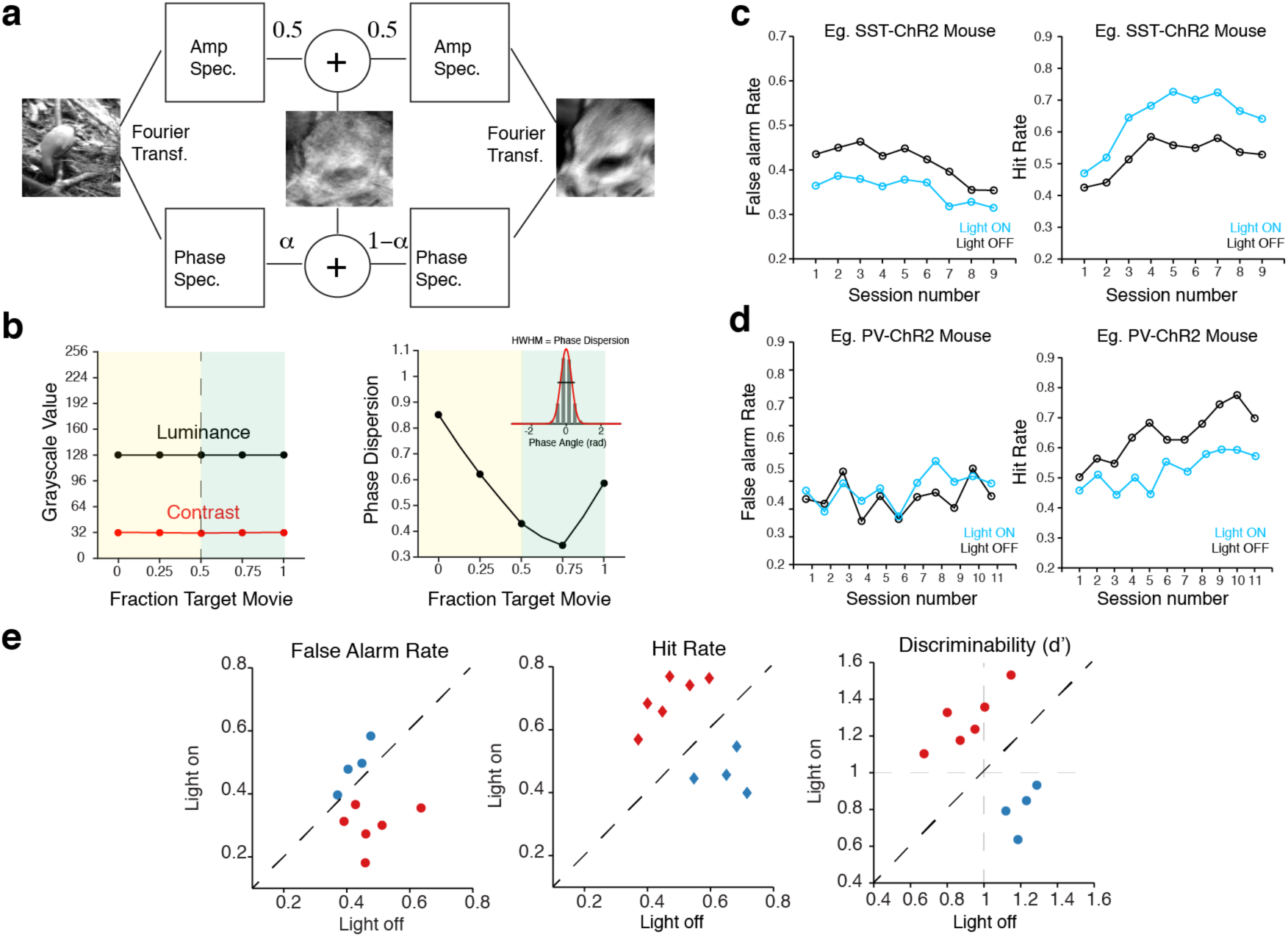
Description of phase-randomized movies and the effect of optogenetic manipulation on hit rate, false alarm rate and discriminability. **(a)** Schematic illustrating of the method used to generate phase randomized movies. Further details are provided in the **Methods**. See also Supplementary Video 2. **(b)** *Left:* Quantification of mean luminance and contrast (standard deviation) of each movie frame in the phase randomized movies. Notice that all movies have been adjusted to have the same mean luminance and contrast. *Right:* Quantification of the phase content of each movie. Unlike the amplitude spectrum, there currently exists no method of parametrically analyzing the phase spectra of natural movies. To compare the distribution of phase angles in each movie, we fit a Gaussian to the histogram of phase angles (see inset). We define the phase dispersion as the half-width-at-half-maximum (HWHM) of this Gaussian fit. Movies with larger phase dispersion have more complex phase spectra with a lot more edges at different orientations. **(c-d)** Representative example from a SST-ChR2 **(c)** and a PV-ChR2 **(d)** mouse showing change in false alarm (FA) rate (lick during non-target movie) and hit rate (lick during target movie) over successive sessions. Each session consisted of approximately 400 trials. On average, both FA and Hit rate remained stable over sessions. **(e)** Scatter plots showing effect of optogenetic stimulation on FA rate (*left*), Hit rate (*middle*) and discriminability (*right*). Data shown is from 4 PV-ChR2 mice and 6 SST-ChR2 mice. PV activation resulted in no significant decrease in FA rate (p = 0.345) but a significant decrease in Hit rate (p = 0.021) and discriminability (p < 0.001). In contrast, SST activation led to a significant decrease in FA rate (p = 0.018), an increase in Hit rate (p < 0.001), and an increase in discriminability (p < 0.001). All P-values computed with Wilcoxon rank-sum test relative to *Light off* condition.

**Supplementary Table 1.** Summary of mouse genotype and viruses used for each experiment sorted by figure number.

| Figure Number | Mouse Used | Virus used |
| --- | --- | --- |
| 1 and S1 | PV-Cre and SST-Cre x Ai14. | AAV1.Syn.GCaMP6f.WPRE.SV40 |
| 1 | PV-Cre and SST-Cre | AAV1.Syn.Flex.GCaMP6f.WPRE.SV40 |
| 2 and S2 | PV-Flp x SST-Cre | AAV1.Syn.fDIO.GCaMP6f.WPRE.SV40 and<br>AAV1.Syn.Flex.NES.jRGECO1a.WPRE.SV40<br>(1:2 mixture) |
| S3a | PV-Flp xSST-Cre | AAV-CAG-Flex-ArchT-tdTomato and<br>AAV1.Syn.fDIO.GCaMP6f.WPRE.SV40<br>(1:2 mixture) |
| S3b | PV-Flp x SST-Cre | AAV1-EF1-dflox-hChR2(H134R)-<br>mCherry.WPRE,<br>AAV1.Syn.Flex.NES.jRGECO1a.WPRE.SV40,<br>AAV1.Syn.fDIO.GCaMP6f.WPRE.SV40<br>(1:1:1 mixture) |
| S4 | PV-Cre and SST-Cre | AAV1.Syn.GCaMP6f.WPRE.SV40<br>AAV2/1-HSYN-DIO-hM3Dq-mcherry (or<br>AAV2/1-HSYN-DIO-hM4Di-mcherry)<br>(1:1 mixture) |
| 3 | PV-Cre x Ai32 | AAV1.Syn.GCaMP6f.WPRE.SV40 and<br>AAV2-CAG-FLEX-tdTomato (1:2 mixture) |
| 3 | PV-Flp x SST-Cre | AAV1-EF1-fDIO-hChR2(H134R)-<br>mCherry.WPRE<br>AAV1.Syn.Flex.GCaMP6f.WPRE.SV40<br>(1:2 mixture) |
| 4 | SST-Cre x Ai32 | AAV1.Syn.GCaMP6f.WPRE.SV40 and<br>AAV2-CAG-FLEX-tdTomato (1:2 mixture) |
| 4 | PV-Flp x SST-Cre | AAV1-EF1-dflox-hChR2(H134R)-<br>mCherry.WPRE,<br>AAV1.Syn.fDIO.GCaMP6f.WPRE.SV40<br>(1:2 mixture) |
| S5 | PV-Cre and SST-Cre x Ai32 | AAV1.Syn.GCaMP6f.WPRE.SV40 and<br>AAV2-CAG-FLEX-tdTomato (1:2 mixture) |
| S6 | PV-Cre and SST-Cre x Ai14 | AAV1.Syn.GCaMP6f.WPRE.SV40 |
| 6 | PV-Cre x Ai35 | AAV1.Syn.GCaMP6f.WPRE.SV40 and<br>AAV2-CAG-FLEX-tdTomato (1:2 mixture) |
| 7 | PV-Cre and SST-Cre x Ai32 | No virus |

**Supplementary Table 2.** Summary of training stages. Mice have to achieve at least 70% correct on each stage before moving to the next stage.

| **Stages** | **Target movie Probability** | **Stimuli order** | **Reward Type** | **Reward available (from movie start, s)** | **Number of Sessions (Ave.)** |
| --- | --- | --- | --- | --- | --- |
| 0 | 40% | Deterministic | Auto | 0.5 | 10 |
| 1 | 20% | Deterministic | Auto | 0.5 | 3 |
| 2 | 20% | Deterministic | Lick to initiate | 0.5 | 5 |
| 3 | 20% | Deterministic | Lick to initiate | 0.7 | 3 |
| 4 | 20% | Random | Lick to initiate | 1.0 | 8 |

## METHODS

### Experimental Animals

All experiments were carried out under protocols approved by MIT’s Committee on Animal Care and conformed to NIH guidelines. The main mouse lines used in this study are: Pvalb-IRES-Cre (PV-Cre, Jax: 008069), Sst-IRES-Cre (SST-Cre, Jax: 013044), Pvalb-2A-FlpO-D (PV-Flp, Jax: 022730), Ai32 (RCL-ChR2(H134R)/EYFP, Jax: 012569), Ai35 (RCL-Arch/GFP, Jax: 012735) and Ai14 (RCl-TdT-D, Jax:007908). All mice (see Supplementary Table 1) were maintained on a C57BL6/J background. Only mice older than 8 weeks old were used in this study. Mice were housed in the vivarium on a standard 12 hour light/dark cycle and were housed at maximum 5 mice in each cage. Experiments were performed during the light portion of the cycle.

To create SXP mice, we crossed homozygous male SST-Cre mice with heterozygous female PV-Flp mice. Pups from the first off-spring generation (F1) were genotyped at postnatal day 21 using a commercial service (Transnetyx), and only pups that expressed Cre and Flp were selected. Only F1 pups from 3 breeder lines were used in this study.

To create PV-ChR2, SST-ChR2 and PV-Arch mice, we crossed homozygous male PV-Cre or SST-Cre with heterozygous female Ai32 or Ai35 mice. Again, F1 pups were genotyped at postnatal day 21 using a commercial service (Transnetyx), and only pups that expressed Cre and GFP, the fluorophore attached to ChR2, (or EYFP in the case of Ai35 mice) were selected. Only F1 pups from 6 breeder lines were used in this study.

### Surgical procedures

Adult mice (between 8-10 weeks old) were an anesthetized with 1–2% isoflurane (vol/vol) and a sterile surgery was performed described previously ^33,64^. First, a small circular piece of scalp was excised to expose the skull. After cleaning and drying the skull using a razor blade and sterile cotton swabs, a custom-built head-post was implanted to the exposed skull with cyanoacrylate glue (Loctite) and cemented with dental acrylic mixed with black paint (C&B Metabond). A craniotomy (3 mm in diameter) was made over the left V1 (2.5 mm lateral and 0.5 mm anterior to lambda). Care was taken not to damage the dura during the craniotomy.

Depending on the experiment, a cocktail of adeno-associated viruses (AAVs, described in list below) were then injected using a beveled pipette (20-30-μm diameter tip Drummond Scientific) backfilled with mineral oil at a speed of 50 nl/min at 5-6 injections sites. Between 100-150 nl of virus was injected per injection site. After each injection, pipettes were left in the brain for an additional 5-10 minutes (depending on injection volume) to prevent backflow and to ensure proper virus spread. Following virus injections, a chronic imaging window was placed in the craniotomy. The imaging window consisted of an inner 3mm glass window and an outer 5mm glass window (Warner Scientific), which were glued together using optically transparent UV curing glue (Norland Optical). Once mice recovered from anesthesia, they were returned to their home cage and were singly housed. Mice were provided with analgesia (meloxicam, 0.1 mg per kg of body weight) subcutaneously three days post-surgery. Imaging experiments typically started 14 to 21 days post-surgery to allow for sufficient viral expression and recovery. Mice with limited optical access due to bone growth or infection were excluded from further analysis.

### Two-photon imaging

Imaging was performed using a Prairie Ultima two-photon system (Bruker) driven by two Spectra Physics Mai-Tai lasers, both passed through a Deep-See modules (Spectra Physics). Imaging was performed using high performance objective lens (Olympus XL 25x Plan N objective, NA = 1.05). In most experiments (except dual wavelength imaging, see below), we tuned the laser to 965nm to enable us to optimally visualize both GCaMP6f and tdTomato fluorescence. To separate red and green fluorescence, we used a 565nm dichroic filter, a 520/40nm green filter and a 600/50nm red filter (all from Chroma). We used a removable curtain made from blackout material (Thorlabs) and a custom holder to isolate the visual display from the microscope.

In dual wavelength imaging experiments, we tuned one laser to 920nm to excite GCaMP6f and another laser to 1020nm, which was the limit of the laser, to excite jRGECO1a. Both laser beams were multiplexed using a half wave plate and a polarizing beam splitter (Thorlabs) before being focused onto a pair of galvanometer mirrors. In doing so, mirrors scanned both laser beams over the same neural field-of-view. This allowed us to image from jRGeco1a-expressing SST and GCaMP6f-expressing PV neurons within the same neural population simultaneously.

In all experiments, images were acquired using ScanImage3.8 in Matlab (Vidrio Technologies) at 20 Hz, 512 × 100 pixels (2x optical zoom). The images covered a cortical area of approximately 150 μm × 150 μm. Images were collected at a depth of 180-280 μm below the pial surface, which corresponds to cortical layer 2/3. Prior to imaging, mice were habituated to head-fixation for 2-3 sessions to reduce stress and anxiety. Typically, 5-8 non-overlapping fields-of-view (FOV, each an independent neural population) was collected for each mouse. FOVs were determined by hand mapping the receptive field locations of neurons in the FOV by moving a sinusoidal grating within a 20-degree Gabor-patch around the screen in 20x20 degree square patches. FOVs without visually evoked responses to these stimuli or those with receptive fields close to the edges of the monitor were discarded.

### Visual stimuli

Natural movies from Van Hateren database as previously described^33^, were displayed on a gamma-corrected, 7-inch 1080p LCD computer monitor (Xenarc) placed 3 inches in front of the contralateral eye. All movies were in grayscale. This computer monitor covered a visual space of approximately 50x70 degrees. Stimulus timing was controlled using Psychtoolbox-3 with custom written Matlab (Mathworks) scripts. Each movie was presented for 4s (30 frames/s) and were interleaved with 4s iso-luminant gray screen. Each movie frame was adjusted to have a luminance of 128 (mean of pixel histogram) and an RMS contrast of 32 (RMS of pixel histogram) on a 0-255 grayscale using the SHINE toolbox^65^.

### Chemogenetic activation and inactivation

Chronic IN activation and inhibition was accomplished with Cre-dependent DREADD (hM3Dq or hm4D1 respectively) in PV-Cre and SST-Cre mice. AAV viruses encoding these DREADDs were co-injected with an AAV viruses encoding GCaMP6f into layer 2/3 of visual cortex, following which mice were implanted with a cranial window as described above. On the day of the experiment, a field-of-view was first chosen by hand-mapping spatial receptive fields of EXC neurons (mCherry-negative) as described above. We measured the reliability of EXC and INs within this field of view to repeated presentations of natural movies as described above (“before condition”). The spatial location of this field-of-view was noted and reference images were taken. Clozapine-N-Oxide (CNO, Sigma-Aldrich) was dissolved in 0.9% sterile saline to an effective concentration of 1mg/kg. Mice received an intraperitoneal injection of either CNO or 0.9% saline (control) after one imaging session and were allowed to recover in their home cage for approximately one hour before imaging commenced again. We used the references images and the spatial location of the find the same field of view. We were consistently able to find the same population of cells before and after CNO administration.

We took several steps to ensure that the same cells were include in the “before” and “after” conditions. First, we registered both fields-of-view by maximizing cross correlation to the same reference image (CV template matching, ImageJ). Next, the same ROI masks were used to segment images collected after Saline or CNO administration. Cells in the after condition that moved relative to the original ROIs were not analyzed.

### Optical activation and inactivation

A 473nm (blue, 200mW peak power) laser and a 532nm laser (300 mW peak power, both from Opto Engine LLC) were used to activate ChR2 and Arch respectively. Both lasers were coupled to a 0.12 NA optical fiber (Thorlabs) and these fibers were launched into the uncaging beam path of the two-photon microscope. The uncaging beam path was co-aligned with the imaging path such that the single wavelength laser illuminated the same FOV as the two-photon laser. In this way, we were able to provide focused activation (or inactivation) of neurons within the same FOV. These single wavelength lasers were triggered using a TTL pulse generated by the visual stimulus computer (see description below). Laser power at the tip of the objective was 1.5mW for ChR2 and 2.5mW for Arch experiments respectively. Laser power was measured before the start of each experiment. Using single cell imaging, we determined that these laser powers were sufficient to reliably drive activation (or suppression) of PV and SST neurons (see supplementary figure). These power values are also consistent with previously published reports using similar mice^45,66^.

In all experiments, we used a stimulus-triggered random stimulation protocol to activate/inactivate cells. Each stimulation pattern consisted of four 20ms pulses of laser with a 10ms inter-pulse interval (i.e. 110 ms total duration per stimulation epoch). In Arch experiments, we used 2x40ms laser pulses with a 5ms inter-pulse interval (175 ms total duration). This pulse pattern was applied at 22 different frames during a natural movie. The frame numbers that triggered the pulses were fixed for each experiment. Specifically, the first pulse occurred at stimulus onset (i.e. triggered by frame number 1), the last pulse occurred at stimulus offset (frame number 240) and the remaining 20 pulses were chosen at a fixed interval. Before the start of each experiment the order of these pulses were pseudo-randomized such that, in each experiment, all pulses appeared in random to each other. The random order was noted and used for *post hoc* analysis (described in the next section). This was done to minimize spurious network activity caused by rhythmic photo-stimulation. In order to calculate reliability, each pulse pattern was repeated 10 times. This resulted in a total of 220 laser-on events and 10 laser-off events (used for controls). Also, to prevent adaptation to repeated presentations of one movie, we interleaved the “pulsed” movie with a “non-pulsed” movie, during which no laser was applied. As a consequence, the network was allowed at least 8s to recover before the next laser pulse was applied.

To determine which two movies to select, we first presented 40 repetitions of five different movies and computed reliability of each neuron in that FOV as described below. Movies with the highest two reliability values were then selected as the “pulsed” and “non-pulsed” movie respectively. This method was repeated systematically for each FOV, and helped us reduce the number of unreliable or non-visually responsive neurons.

### Visually responsive neurons and spike rate inference

All data analysis was performed with custom written Matlab (Mathworks) and ImageJ (NIH) macros (NIH) that called built-in functions. Following imaging, images stacks (tiff format) were first corrected for motion artifacts using an open-sourced ImageJ plugin (https://sites.google.com/site/qingzongtseng/template-matching-ij-plugin) that maximized the cross-correlation coefficient between frames. Frames with excessive motion artifacts that could not be corrected were also discarded. Frames with photostimulation laser artifacts were also discarded from analysis (usually 1-2 frames) and cubic spline interpolation was used to smooth over these blanked frames.

Next, neuronal ROIs were manually segmented in ImageJ (NIH) using the Cell Magic wand tool (https://www.maxplanckflorida.org/fitzpatricklab/software/cellMagicWand/) and fluorescence time series for each neuron was computed by averaging pixels within each ROI. A modified version of the Cell Magic wand tool was used to identify a surrounding neuropil region, which was an annulus of outer diameter = (15 pixels + diameter of cell) around each cell. This data was then imported into Matlab for further analysis. The raw fluorescence of each cell was computed using the formula: *F* = *F*_*Raw*_ − *F*_*neuropil*_.

All data analysis was performed using custom written scripts in Matlab. Significantly visually responsive cells were determined from the fluorescence time changes (ΔF/F) by performing a one-tailed Student’s t-test between visually evoked (4s movie on) and spontaneous responses (4s gray screen in between movies). To obtain a better estimate of the spontaneous activity we also collected 120 s of activity to an iso-luminance gray screen before the start of each experiment. Only cells with p < 0.001 were classified as visually-responsive.

We used a two-step procedure to estimate the firing rates of visually-responsive cells. We first detected statistically significant calcium transients from the ΔF/F time series of each neuron by analyzing the distribution of positive-going and negative going calcium transients as described previously (Dombeck et al., 2007; Danielson et al., 2016). This method allowed us to minimize the number of false-positive calcium transients induced by brain motion (false-positive error rate <1%). Next, we filtered the ΔF/F time series of each neuron such that non-significant transients were 0, while significant transients were untouched. Following this, we used an optimized deconvolution algorithm for GCaMP6f^67^ to infer the firing rate for each neuron. Briefly, this algorithm inferred the probability of spiking from statistically significant calcium transients. To convert this probability into a firing rate (measured in events/s), we multiplied each probability by 20 Hz, the frequency at which the calcium transients were sampled. Unless otherwise stated, all data analysis was performed using inferred firing rates.

### Change in firing rate and change in reliability following photostimulation

In all photostimulation experiments, analysis was restricted to 600 ms time window (12 imaging frames) following laser activation. Within this time window, we determined the change in firing rate using the following formula

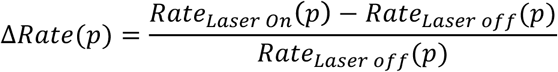

where *p* is the pulse number. Since each neuron responded at different time points of the movie, averaging this across population of neurons would obscure any changes in the firing rate (or reliability). Thus, we aligned the firing rate trace of each neuron obtained on the Laser-off trials (control condition) such that the maximum rate occurred at 1s following stimulus onset. This index was then used to align the firing rate traces on the Laser-on trials.

Response reliability to natural movies was calculated using the equation:

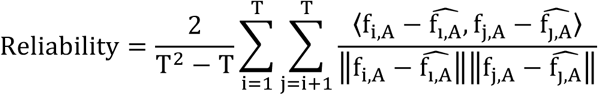

where i,j ∈ [1,T] index trial numbers and f_i,A_ is the rate on the i^th^ trial for movie A, 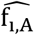 is the average rate for that trial and 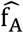 is the average across trials (mean rate). Thus, from this equation the response reliability is the average correlation of all pairwise combinations of trials, corrected for differences in mean firing rate^33^. Similarly, we computed an unbiased estimate of the firing rate variance between trials for movie A using the formula

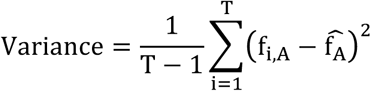

In photo-stimulation equations, the change in reliability induced by laser activation was calculated using the formula:

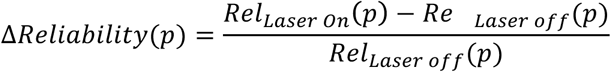

Similar to the firing rate, we aligned the reliability on the Laser-off trials such that each neuron was maximally reliable at 1s. The same time index was then used to align reliability on the Laser-on trials. To determine the epoch of maximum and minimum reliability, we calculated the time index corresponding the maximum and minimum reliability on the Laser-off trials from unaligned traces. For the regression analysis shown in Figures 3, 4 and 6, we computed the change in variance using the simple formula,

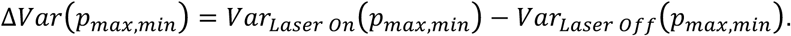

A similar formula was used to calculate Δ*Rate* and Δ*Reliability*.

### Behavior: Water restriction and Rig

PV-Cre x Ai32 (PV-ChR2) and SST-Cre x Ai32 (SST-ChR2) mice were implanted with a cranial window and a head post. Mice were allowed to recover from surgery for 1-2 weeks before beginning water restriction. Specifically mice were placed on a water restriction schedule in which they received a minimum daily amount (40μL water per gram, daily mouse weight) ^68^. Weight, behavior and general condition, were monitored by veterinary staff.

Mice were trained on a custom-built behavior rig ^64^. Visual stimuli were presented on the same Xenarc monitors used in physiological experiments (described above). In this set-up the monitors also covered a visual space of ~70x90 degrees. Water was delivered to the mice via a conductive lick spout ^68^.

### Behavior: Training schedule

Once stable body weight was reached, mice began training on the natural movie discrimination task. Typically, mice were trained daily (7 days a week) in one-hour sessions. On the first two-three training days, mice were head-fixed and given water reward in order to acclimatize to both head fixation and drinking from a waterspout. The two movies that evoked the most reliable responses were selected to be the target and non-target movies respectively. Each movie was presented for 2s (30 frames/s) and were interleaved with 2s isoluminance gray screens. We presented these movies using an “odd-ball” paradigm, such that the target movie appeared with a lower probability than the non-target movie. In early stages of training (see below), the target movie appeared with a probability of 20% and were presented in deterministic order, such that one target movie appeared after a run of four non-target movies. In later stages, we randomized the order, such that the mouse could not use movie history to predict the next movie. We reasoned that this “odd-ball” paradigm would keep the mice more engaged in the task as the relevant stimuli only appeared rarely.

A summary of the different training stages is shown in Supplementary Table 2. In the first stage (Stage 0), mice were taught how to associate a water reward with the target movie. Specifically a 5μL water reward was given to mice 2s into the target movie. Once mice achieved a stable hit rate (HR) of 90% over three consecutive sessions, mice were graduated to Stage 1. In Stage1, the non-target movie was introduced (80% of trials) and mice were trained to withhold licking. False alarms (FA) were indicated with a 200ms white noise burst. In this stage, mice were auto-rewarded on every target movie trial. Once correct reject rate (CR) reached 80% mice were graduated to the next stage. In Stages 2-3 the probability of auto-reward was gradually reduced from 50% (Stage 2) to 25% (Stage3). Mice were promoted to Stage 4 only once they were able to achieve a CR and HR of 80%. In Stage 4, target and non-target movies were played in a deterministic sequence but no auto-reward was given. Finally in Stage 5, mice were trained on the randomized sequence. Once mice reached a HR and CR > 70% on the randomized sequence they were tested on the Target discrimination version of the task. Typically, training lasted for 3-4 weeks with mice completing 400-600 trials per day.

### Behavior: Movies used in classification task

In this version of the task, three more movies were introduced and the mice had to determine which movies from this ensemble had more target-like features. Specifically, to create these movies we blended the phase spectrum of the non-target movie with the phase spectrum of the target movie according the formula

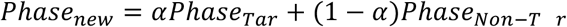

with *α* = [0,0.25,0.5,0.75,1]. Also we replaced the amplitude spectrum of all movies with the mean amplitude spectrum of the non-target and target movie.

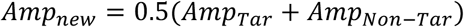

All movies were corrected to have the same mean luminance and contrast using the SHINE toolbox ^65^. Movies with *α* = [0.75,1] were treated as target movies and presented with a probability of 0.4, whereas movies with *α* = [0,0.25,0.5] were treated as non-target movies and were presented with a probability of 0.6. Non-target movies were not rewarded except for *α* = 0.5, which was rewarded on 50% of the trials. On average mice completed between 400-800 trials of the classification task per day.

We compared similarity between the phase-blended stimuli with the target movie using the structural similarity index (SSIM). The SSIM uses image structural information, such as mean, variance and covariance, to estimate dependencies between pixels (Wang et al., 2004). Specifically, we computed SSIM between images *i* and *j* using the following equation:

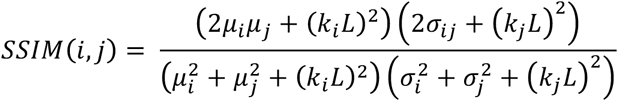

where, *μ*_*i,j*_ and *σ*_*i,j*_ is the mean and standard deviation of images *i* and *j* respectively, *σ*_*i,j*_ is the covariance and *L* is the dynamic range of the image. For further details, see (Wang et al., 2004).

### Behavior: Behavioral testing and optogenetic stimulation

Once mice were proficient at this task, we optogenetically activated ChR2-expressing neurons using a fiber-coupled 470nm LED source on 50% of the trials. Here, we used 8 pulses, 10ms each with a 10ms inter-pulse-interval (150 ms in total). The onset of stimulation was coincident with the movie onset. The average power at the tip of the fiber was 8mW, and the power spectral density was similar to what we used for physiology experiments. We assessed performance by computing the hit rate (HR) and false alarm rate (FAR) using the following equations:

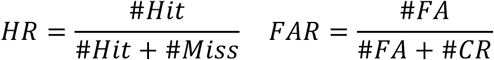

where #*Hit* is the number of licks and #*Miss* are the number of misses to the target-biased movies, and #*FA* is the number of licks and #*CR* number of misses to the non-target-biased movies Using these equations, we computed the discriminability (*d*′) between target and non-target movie (Figure S9) as,

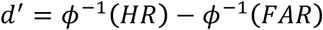

where *ϕ*^−1^ is the CDF of a standard Gaussian with 0 mean and 1 variance, which was computed using the *norminv* function in Matlab.

To determine the effect that LED activation had on task performance, we fit sigmoidal psychometric functions to the proportion of correct response independent on LED-on and LED-off trials using the following equation:

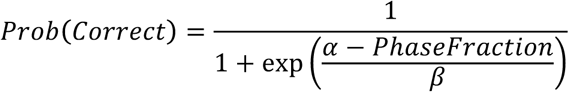

where parameters *α* and *β*, which represent the bias and slope respectively, were determined using least squares regression. Specifically, increasing *β* decreases the slope indicating less sensitivity to the phase fraction. Increasing *α* shifts the psychometric to the right, indicating an increase in detection threshold. Thus, an improved performance (more sensitive at detecting the target movie) at this movie classification task manifests in a reduction both in *α* and *β*. Only trials with fit quality >85% were kept (12/18 session). Significance of the fit was determined by performing a chi-squared test. All changes were compared to *LED-off* trials.

### Multi-unit rate-based neural network model

We built a four-unit rate-based model of layer 2/3 of visual cortex to study the effects that the SST-PV dynamics had on EXC neuron reliability. Each population was represented by a single rate-based equation:

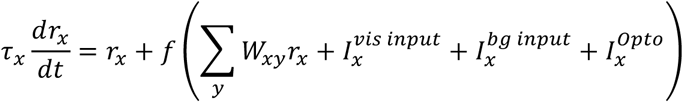

where *r*_*x*_ is the firing rate of the cell population *x* (EXC, PV, SST, VIP). PV units had a time constant *τ*_*x*_ = 10*ms*, while EXC, SST and VIP units had a slower time constant *τ*_*x*_ = 20*ms*. We modeled the rate-current transfer function of each population using the power-law function:

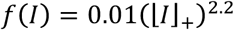

EXC, PV and SST units received “visual input” 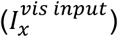 which accurately reflected the temporal activity of natural movie stimulation (Supplementary Fig. 8a, b). This visual input was the summed activity of a bank of 50 linear-nonlinear-Poisson units. The linear filter consisted of a spatial log-Gabor receptive field (total 6 different orientations (0-180°) and ranged in size from 12-18° of visual angle) and gamma functions with a range of temporal delays (140-200ms). These spatiotemporal receptive fields closely resemble those seen in mouse visual cortex ^33,69^. Because we did not know the locations of the RFs *a priori*, we randomly picked 50 possible locations on the screen. The same natural movies used in our experiments were first convolved with each spatio-temporal log-Gabor filter, which was then rectified with a point-wise nonlinearity to produce a firing rate estimate. This firing rate estimate was then used to generate an inhomogeneous Poisson spike train. To generate input to the different units in the model, we filtered this Poisson spike train through a facilitating alpha synapse to generate the current 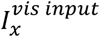. The weight of each synapse was varied from trial-to-trial, which together with the stochastic nature of the Poisson process, created trial-to-trial variability in the model that closely resembled the variability observed between movies (Supplementary Fig. 8c). EXC and PV units received summed input from units with smaller log-Gabor sizes while SST unit received input from larger log-Gabor sizes reflecting differences in preferred spatial frequencies of each cell type^36^. VIP units did not receive any visual input.

All units received background current 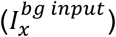, which was modelled as a stochastic Poisson process. We varied input rates between the different units (EXC and SST = 10 Hz, PV = 24Hz, VIP =15 Hz) to match spontaneous firing rates observed *in vivo*. In this way, the spontaneous activity was uncorrelated between each neuronal subtype in our model, and therefore represented an independent source of noise.

In experiments with optogenetic perturbation, we modelled optogenetic input into both PV and SST units 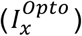 as a train of square wave pulses which mimicked the temporal properties the laser stimulation used in our experiments. To test for robustness and to mimic natural trial-to-trial variability of ChR2 and Arch, we varied the pulse amplitude by +/- 10% in each trial by drawing values from a uniform distribution. The amplitude values were: SST-ChR2 = 25 mA, PV-ChR2 = 45 mA, PV-Arch = −50 mA.

Connections between populations were given via the following weight matrix:

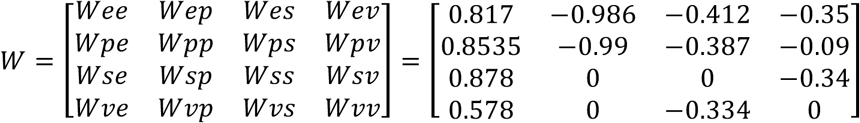

where *W*_*xy*_ is the weight of the connection from neuron *y* to neuron *x*. In some experiments, we removed the SST→PV connection by setting *W*_*PS*_ = 0. This weight matrix reflects the known connectivity between neuron subtypes in the visual cortex, and is derived from previously published results^39,70^.

To compute reliability, we simulated 30 trials with the same visual stimulus, and used the same equation defined above to compute EXC unit reliability. To test the robustness of our model to parameter changes, we created 500 models (each dot in Fig. 5) by independently varying all the parameters of the model by +/- 10% of their current values. Numerical integration was performed in Matlab using the forward Euler method with a time step of 0.05 ms.

To determine which factors (e.g. PV suppression, EXC suppression, etc.) contributed the most to the changes in EXC unit reliability following PV/SST activation/suppression, we performed multivariate linear regression using the following model:

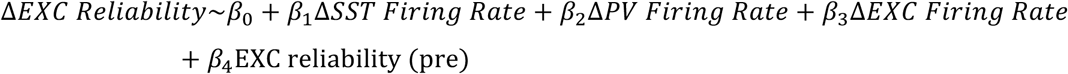

Linear regression was performed in Matlab (*fitlm*) and statistical tests (Students T statistic) were computed to assess the significance of each predictor.

### Statistical Analysis

All statistical analysis was performed using custom written scripts in Matlab and R. No tests were conducted to determine sample size. Data were first tested for normality using the Shapiro-Wilk Test. All data presented in this paper are non-normally distributed, thus all statistical tests were conducted using non-parametric statistics. Our experiments involved testing the influence of laser activation on the same population of neurons, thus all comparisons were performed using non-parametric repeated-measures ANOVA (Friedman Test) with Bonferroni’s correction and rank-sum post-hoc tests with significance value was set to 0.05. Post-hoc tests were performed using the two-tailed Wilcoxon rank-sum test relative to the Laser-off condition. To determine if the change in reliability was significant, we performed permutation tests (corrected for family-wise error rate) where we resampled with replacement (10,000 permutations) from the change distribution and tested if the sampled distribution was significantly different from 0 using a one-tailed rank-sum test. Unless otherwise stated, data are presented as median ± 95% CI (calculated using bootstrap sampling). All confidence intervals were determined using bootstrap. All box-whisker plots show median (notch), inter-quartile range (box edges) and data range (whiskers). All p-values are labeled in the figures and their corresponding legends.

### Data and code availability

All scripts used in analysis and model simulations, and raw imaging and behavior data is available from the corresponding author upon reasonable request.

## FOOTNOTES

### Author Contributions

Conceptualization, RVR and MS. Methodology and Resources, RVR, MH and MY. Investigation and Formal Analysis, RVR. Writing, RVR and MS. Supervision and Funding Acquisition, MS.

## Acknowledgements

The authors thank S El-Boustani, M. Andermann, and Sur lab members for helpful discussions; T. Emery and L. Gunter for technical assistance and animal husbandry; R. Neve for creating the FRT-GCaMP6f and FRT-ChR2 constructs; and the Genie Project at the Janelia Research Campus, Howard Hughes Medical Institute for use of GCaMP6f and jRGECO1a. RVR is supported by the HHMI International Student Research Fellowship. This work is supported by NIH grants EY007023 and NS090473 and NSF grant EF1451125, and the Picower Institute Innovation Fund, to MS.

